# Four neurons pattern brain-wide developmental activity through neuropeptide signaling

**DOI:** 10.1101/2025.06.26.661770

**Authors:** Jun Reichl, Julia M. Miller, Harpreet Randhawa, Lesly M. Palacios Castillo, Orkun Akin

## Abstract

The developing brain becomes electrically active before it is ready to process sensory input. During neural circuit maturation, developmental activity is thought to refine synaptic connections by driving neuronal co-activation in rhythmic patterns. The source of this brainwide activity and the mechanisms which regulate its patterns are not well understood. Here we describe cellular interactions that shape developmental activity and their molecular basis. In *Drosophila*, patterned stimulus independent neural activity (PSINA) engages the entire brain in highly stereotyped, globally coordinated cycles of activity. A molecularly-defined population of ∼2,000 neurons (Transient Receptor Potential Gamma, *Trpγ+* neurons) act as an activity template for PSINA. We show that this activity template is patterned by four neurons expressing the neuropeptide SIFamide (SIFa). Signaling through the SIFa Receptor, SIFa modulates the activity of both SIFa and *Trpγ+* neurons to establish the brainwide activity cycles of PSINA. In turn, *Trpγ+* neurons regulate SIFa neuron activity through a recurrent interaction. Neuropeptides act through synapse-free, or wireless, signaling; a fitting mode of communication for a process tasked with refining on-going synapse formation. By placing neuropeptide signaling at the core of developmental activity, this work highlights the rich neurophysiological potential of the chemical connectome in shaping the developing brain.

## INTRODUCTION

At the onset of synaptogenesis, before the nervous system is ready to process sensory input, the developing brain exhibits waves of patterned rhythmic activity. Such developmental activity occurs in many parts of the mammalian brain, including the visual, auditory, and olfactory systems, the spinal cord, thalamus, and the cortex^1–3^. Altering the activity in sensory systems, where it has been studied in greatest depth, leads to disruptions in topographic mapping^4–6^ and circuit function^7,8^. Changes to the activity in the developing cortex are thought to be linked to neurodevelopmental disorders^2^, highlighting the significance of understanding how this process contributes to the assembly of the brain. While specifics are outstanding, and likely to be case-specific, the consensus view is that the function of the activity is to mediate neuronal co-activation^9^: Selective strengthening or weakening of synapses through Hebbian plasticity^10^. This framework elevates the importance of the spatiotemporal patterns of activity^11^ and, by extension, the mechanisms which generate them.

For the few cases where molecular and cellular factors necessary for wildtype activity patterns have been identified^12–14^, they have proved to be valuable handles on the biology of developmental activity. The best studied example is the so called ‘retinal waves’^15,16^ which initiate in the developing retina and spread to other visual areas, including the superior colliculus and the visual cortex^17^. In rodents, retinal waves appear two days before birth and persist for two weeks, until eye opening. For much of this time, the waves depend on cholinergic signaling from the network of starburst amacrine cells (SACs), both to each other and to retinal ganglion cells (RGCs)^12^. Loss of the β2 subunit of the nicotinic acetylcholine receptor, which mediates this cholinergic communication, severely alters retinal waves^18^ and leads to RGC projection defects in higher visual areas^19,20^. While network interactions among SACs are thought to be sufficient to initiate retinal waves^12^, coordination of the activity between the two eyes^17^ indicates that there are additional regulatory mechanisms which shape the patterns. Notably, how the core mechanisms that initiate developmental activity are regulated to give rise to the final patterns is not well understood in any part of the developing brain. A promising candidate for bi-lateral retinal wave coordination is a subtype of RGCs which project to the contralateral retinas^21^. In addition, brain-to-retina projecting histaminergic neurons from the hypothalamus have been shown to modulate visual responses in mice^22^, suggesting that regulation need not arise autonomously in the eyes. The potential for such long-range activity modulation was highlighted by a recent report showing that hypothalamic peptidergic neurons project into nearly all areas of the mouse brain^23^.

The discovery of developmental activity in the fruit fly *Drosophila melanogaster*^24,25^ indicates that this phenomenon fundamental to neurodevelopment. The adult fly brain is built during the 100 hours of metamorphosis, or pupal development. Patterned, stimulus-independent neural activity (PSINA, *‘see-nah’*) starts ∼50 hours after puparium formation (hAPF)^25^, co-incident with the appearance of the first structural synapses^26,27^. For the following two days until an hour before the adult fly emerges, PSINA engages the whole brain in cycles of activity comprising globally coordinated active and silent phases (Fig.1a-c). Embedded within each global active phase are stereotyped cell-type-specific activity patterns which are more correlated between future synaptic partners^25^.

**Figure 1.**
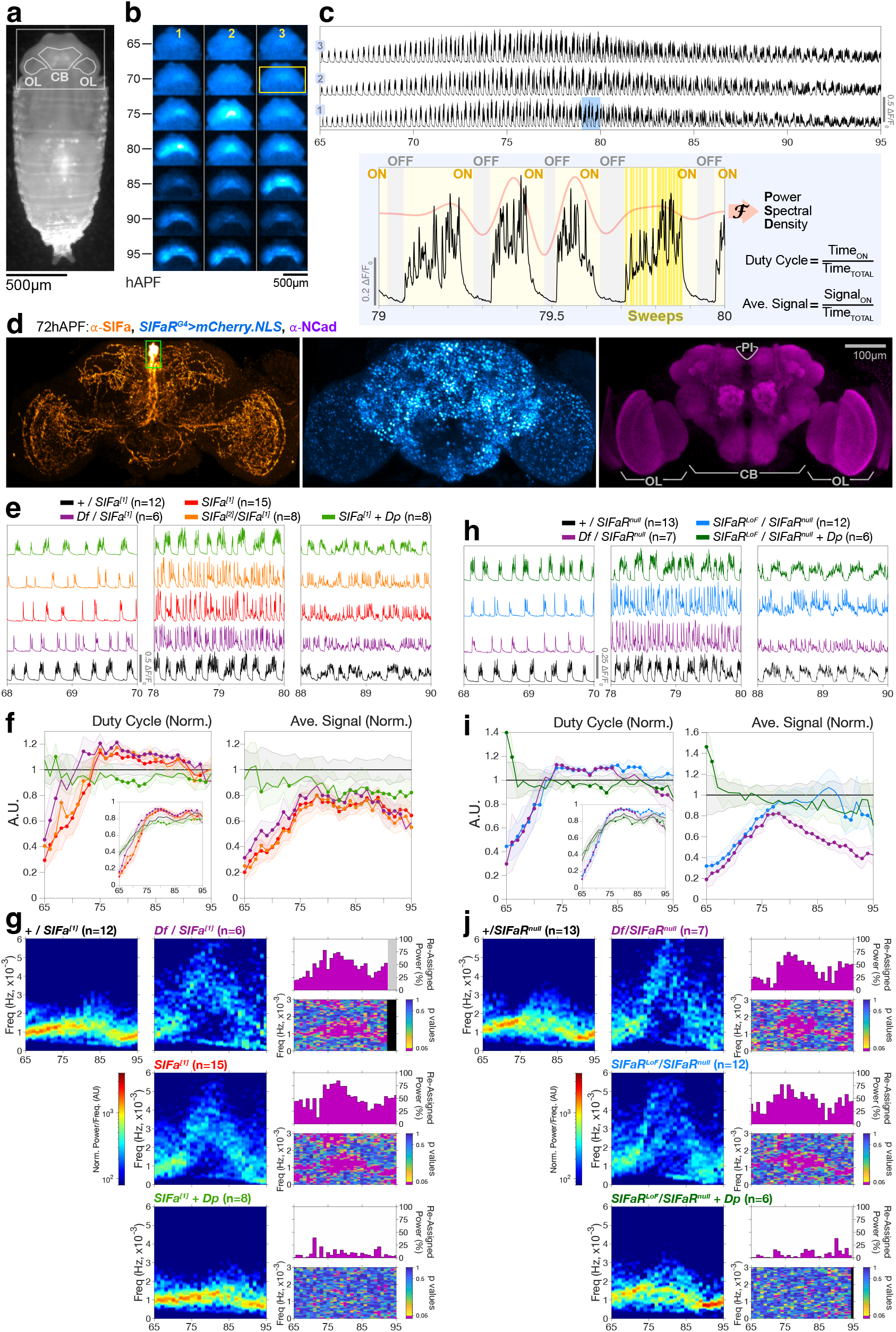
*SIFa* and *SIFaR* are necessary for PSINA cycles. **a**. 65 hAPF *D. melanogaster* pupa. Cuticle around the head (boxed) has been removed for widefield imaging. Brain is outlined: CB, central brain; OL, optic lobe. Scale bar, 500μm. **b**. Montage of pupal heads from one widefield imaging session. Signal is fluorescence from pan-neuronally expressed GCaMP6s. Yellow rectangle (top-right) is typical region-of-interest (ROI) used to extract PSINA traces. All time axes in this and following figures are in units of hAPF. Scale bar, 500μm. **c**. *Top:* Wildtype PSINA recorded from enumerated pupae in (b); sampling rate is 0.5Hz. *Bottom:* Expanded view of PSINA from the blue highlighted 1h span above depicting the active (ON) and silent (OFF) phases of activity cycles, sweeps, and principal metrics used in analyses. PSD, power spectral density: PSINA traces were filtered to remove sweeps (coral overlaid profile), and PSD was calculated using the Discrete Fourier Transform (‘F’). **d**. Maximum intensity projection (MIP) of 72hAPF brain with *SIFaR*^*G4*^ driving *mCherry.NLS* (native fluorescence, cyan-hot look-up table (LUT)); brain also stained against SIFa (orange-hot LUT) and N-Cadherin (magenta). Green rectangle boxes SIFa soma. CB, central brain; OL, optic lobe; PI, Pars Intercerebralis. Scale bar, 100μm. **e-g**. *SIFa* mutant analysis **e**. Representative PSINA traces, color-matched to genotypes shown; numbers are sample sized used in (f) and (g). *Df*, deficiency; *Dp*, genomic duplication. **f**. Duty cycle and average signal, normalized to control genotype (black). Inset in duty cycle plots true values of metric. Shaded areas, standard deviation. Closed circles mark where experimental distribution differs from control (p<0.05) by two-sample KS test. **g**. Average PSDs from control (first column) and experimental (second column) animals. PSD for each PSINA trace was calculated at every hour within a 2-h sliding window and normalized to remove contribution from changes in signal amplitude. Displayed PSDs are averages of replicates for each condition. Labels color-matched to (e,f); sample sizes indicated. PSD comparison of *SIFa*^*[1]*^*/SIFa*^*[2]*^ to +/*SIFa*^*[1]*^ omitted for brevity. In third column, bottom panel of each set maps p values from KS tests of control and experimental PSD distributions at each frequency-time coordinate; p<0.05 displayed in magenta. Top panel is fraction of control signal power re-assigned to different (p<0.05) frequencies in the experimental data. See also Extended Data Fig.1. **h-j**. *SIFaR* mutant analysis with *SIFaR*^*LoF*^ (i.e., *SIFaR*^*GFSTF*^); data presented as in (e-g).

PSINA relies on a population of ∼2,000 neurons, genetically defined by their expression of the cation channel Trpγ^28^. Loss of *trpy* gene function alters PSINA patterns both across the brain and at the single cell-type-level and disrupts synaptic development in a non-cell autonomous manner. Brainwide PSINA in *trpγ* mutants mirrors the activity recorded from *Trpγ+* neurons^28^, indicating that this small population acts as an activity template for the rest of the brain.

Here, we report that the *Trpγ+* activity template is modulated by the neuropeptide SIFa. SIFa is a highly conserved^29^ arthropod neuropeptide produced principally by four neurons in the Pars Intercerebralis (PI) region^30,31^(Fig.1d), a neuroendocrine command center homologous to the mammalian hypothalamus^32^. SIFa neurons (aka DNc01/2^33^) are present in the larva and undergo extensive remodeling during pupal development to extend processes throughout the adult brain and ventral nerve cord^30^. The SIFa Receptor, SIFaR^34^, is a G protein-coupled receptor (GPCR) expressed broadly throughout the brain (Fig.1d). In the adult, SIFa-SIFaR signaling has been implicated in a range of adult behaviors, including feeding^35^, activity rhythms^36^, sleep^37,38^, and courtship^30,39^. We find that, during pupal development, this signaling module acts on *Trpγ+* neurons to pattern the brainwide activity cycles of PSINA. By identifying four defined neurons that shape brainwide activity through a specific ligand-receptor interaction, our work brings into focus a genetically-programmed neural engine of discrete composition driving this fundamental phase of neurodevelopment.

## RESULTS

To observe PSINA, we use a live imaging approach we term the widefield assay (Supplementary Video 1). We use genetically encoded calcium indicators to visualize neural activity: GCaMP6s^30^ for single cell class recording (e.g., pan-neuronal, PanN, for brainwide PSINA), and GCaMP6s and RCaMP1b^31^ when imaging two different classes of cells. For independent genetic access to different classes of cells, we use the GAL4-UAS^32^ and LexA-LexAop^33^ expression systems. When needed, we control the timing of GAL4-driven expression using GAL80ts^34^, which inhibits GAL4 at 18^°^C, but allows for full expression at 29^°^C. Below, we refer to this approach as temporally-targeted expression. We stage and genotype flies at the onset of pupal development, and, at ∼48hAPF, prepare them for imaging by removing the cuticle around their heads (Fig.1a,b). Using a standard long-working-distance fluorescence microscope, we can record simultaneously from up to 80 pupae for two days; that is, until the flies complete metamorphosis and emerge as adults. This medium-throughput assay made it possible to carry out the neuropeptide screen which identified *SIFa*.

### SIFa and SIFaR are necessary for PSINA cycles

To first assess how total loss of SIFa affects PSINA, we recorded brain-wide activity in whole-animal *SIFa* mutants. Compared to PSINA in controls, *SIFa* mutants show lower signal intensity and altered duty cycle, with less time active per hour during early stages (65-75hAPF) shifting to more time active after ∼75hAPF (Fig.1e,f). Most prominently, the temporal organization of PSINA is altered in SIFa mutants: In the wildtype, PSINA is organized into cycles of active and silent phases with periods of 10-15 minutes (Fig.1c). We visualized these cycles in the frequency domain by taking the Fourier transform of the PSINA traces and calculating the power spectral density (PSD, Extended Data Fig.1); the PSD displays how the power in the original signal is distributed across different frequencies. Most of the power in wildtype PSINA is carried by a band around 10^-3^Hz (Fig.1g, 1^st^ column), which corresponds to the 10-15 minute cycle period. Using the PSD representation, we compared wildtype and experimental (Fig.1g, 2^nd^ column) sets of traces across pupal development and found that the 10^-3^Hz band is absent in *SIFa* mutants (Fig.1g, 3^rd^ column, bottom panels). This change to the PSD is equivalent to ∼50% of the signal power in the wildtype being carried by other frequencies in *SIFa* mutants (Fig.1g, 3^rd^ column, top panels). In effect, the stereotyped activity cycles are lost in *SIFa* mutants. Complementation analysis with a deficiency and rescue with a chromosomal duplication confirmed that the *SIFa* mutant alleles are true nulls and the phenotype is monogenic in origin, respectively (Fig.1e-g, Extended Data Fig.2a). We conclude that *SIFa* is necessary for the stereotyped cycles of PSINA.

We next asked how whole-animal loss of SIFaR, the only known receptor for SIFa^34^, affects PSINA. When paired with a deficiency, the *SIFaR*^*null*^ allele led to the same PSINA phenotypes we observed in SIFa mutants (Fig.1h-j): altered duty cycle, reduced signal, and lost cycles. We tested a protein trap, *SIFaR*^*GFSTF*^, and a gene trap GAL4 driver, *SIFaR*^*G4*^, in combination with *SIFaR*^*null*^ and saw similar changes to PSINA, indicating that these are strong loss-of-function (LoF) alleles (Fig.1h-j, Extended Data Fig.2b-d). Hereafter, we refer to *SIFaR*^*GFSTF*^ as *SIFaR*^*LoF*^. As with *SIFa*, rescue with a chromosomal duplication confirmed the monogenic origin of the *SIFaR* phenotypes (Fig.1h-j, Extended Data Fig.2b-d). We conclude that *SIFaR* is also necessary for the stereotyped cycles of PSINA.

To ask how *SIFa* and *SIFaR* interact, we carried out double mutant analysis using *SIFa[1]* homozygotes and heteroallelic combinations of *SIFaR*^*null*^ with *SIFaR*^*LoF*^ or *SIFaR*^*G4*^ (Extended Data Fig.2e-g,h-j). The *SIFaR* single mutants, particularly those carrying the *SIFaR*^*G4*^ allele, differed from the double mutants in how they affected duty cycle and average signal. We attribute these differences to the *SIFaR*^*LoF*^ and *SIFaR*^*G4*^ alleles not being complete nulls. However, for the most prominent phenotype—loss of cycles—the single mutants were indistinguishable from the doubles. The absence of an additive effect in the double mutants is consistent with SIFa and SIFaR functioning as an exclusive ligand-receptor module. Taken together, our findings indicate that SIFa-SIFaR signaling is required for the stereotyped cycles of PSINA.

### SIFa gene function and SIFa neuronal activity are necessary during PSINA

Analysis of whole-animal mutants does not address when and where *SIFa* and *SIFaR* are required to influence PSINA cycles. To ask whether *SIFa* is necessary during PSINA, we used a GAL4 driver which specifically labels the four large SIFa neurons in the PI^31^ (*SIFa*^*G4* 30^, Extended Data Fig.3a) to knockdown *SIFa* expression in a temporally-targeted manner. The resulting PSINA phenotypes were comparable to that seen in the mutants (Fig.2b,c, Extended Data Fig.3b). We conclude that SIFa expression in the four SIFa neurons during pupal development is necessary for wildtype PSINA.

**Figure 2.**
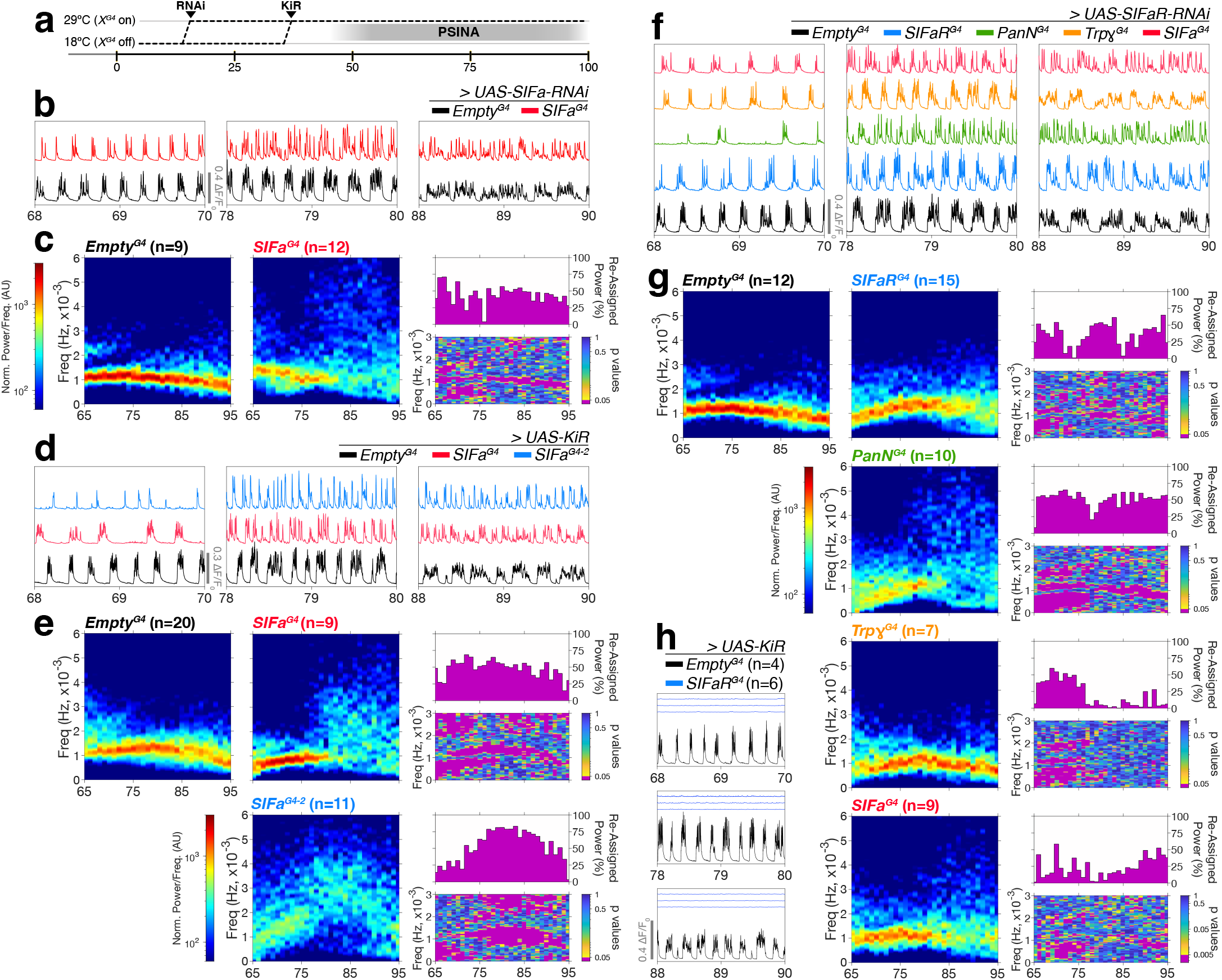
*SIFa* and *SIFaR* are necessary during PSINA. **a**. Temperature schedule used to temporally target RNAi knockdown and KiR2.1 expression to pupal development. **b**,**c**. Temporally-targeted knockdown of *SIFa*. **d**,**e**. PSINA traces, color-matched to genotypes shown. **f**,**g**. Average PSDs from control and experimental animals; presented as in Fig.1g. **d**,**e**. Temporally-targeted KiR2.1 expression in SIFa neurons; presented as in (b,c). **f**,**g**. Temporally-targeted knockdown of *SIFaR*; presented as in (b,c). **h**. Temporally-targeted SIFaR^G4^-driven KiR2.1 expression; one control and three experimental traces shown.

Neuropeptidergic signaling operates alongside co-transmission of other neuroactive compounds^40^. We asked if electrical silencing, by exacerbating the *SIFa* mutant PSINA phenotype, would reveal a co-transmitter requirement in SIFa neurons. Contrary to expectation, *SIFa*^*G4*^ driven, temporally-targeted expression of the hyperpolarizing channel KiR2.1^41^ resulted in a weaker PSINA phenotype (Fig.2d,e, Extended Data Fig.3d). This suggested that the strength of the driver was limiting the severity of the perturbation, so we tested another GAL4 which labels the SIFa neurons, *SIFa*^*G4-2* 42^ (Extended Data Fig.3b). Temporally-targeted KiR2.1 expression by both drivers altered PSINA in similar fashion, and *SIFa*^*G4-2*^ led to a more penetrant phenotype, matching and not exceeding the *SIFa* mutant phenotype (Fig.2d,e, Extended Data Fig.3d). These results are consistent with *SIFa* expression being the dominant contribution of SIFa neurons to PSINA.

### SIFaR gene function during pupal development is necessary and sufficient for PSINA

We pursued a similar strategy to establish the spatiotemporal requirements for SIFaR function. Temporally-targeted knockdown of SIFaR expression with the pan-neuronal *PanN*^*G4*^ driver phenocopied the *SIFaR* mutant phenotype (Fig.2f,g, Extended Data Fig.3e). Knockdown with *SIFaR*^*G4*^ also disrupted PSINA, but was less effective compared to *PanN*^*G4*^, likely due to the dependence of gene trap expression to transcription from the native locus. We conclude that SIFaR gene function is necessary during PSINA.

*SIFaR*^*G4*^ captures an expression domain that expands throughout pupal development to include many thousands of neurons, including the SIFa set (Extended Data Fig.4). This broad expression domain is consistent with a recent fluorescence *in situ* hybridization based characterization of *SIFaR* transcript localization^31^. Notably, we found that temporally-targeted knockdown of SIFaR with *SIFa*^*G4*^ or *Trpy*^*G4*^ also disrupted PSINA (Fig.2f,g, Extended Data Fig.3e). These results suggest that wildtype PSINA relies on both SIFa autocrine signaling in SIFa neurons and paracrine signaling through *Trpy+* neurons. We present further findings on the relationship between SIFa and *Trpy+* neurons below.

In a complementary set of experiments, we asked whether SIFaR expression during PSINA is sufficient to rescue the loss of cycles. Temporally-targeted expression of *UAS-SIFaR*^38^ with *SIFaR*^*G4*^ or *PanN*^*G4*^ in a *SIFaR*^*LoF*^ background led to an overexpression phenotype (Extended Data Fig.5a-d): Beyond restoring the cycles, SIFaR expression from either driver extended the duration of active phases, which also increased the duty cycle and average signal. We were able to match wildtype PSINA more closely by rearing the animals below the fully permissive temperature for GAL80ts, (Extended Data Fig.5e-g) which is expected to reduce the expression levels of the rescue construct. Under both temperature regimes, *SIFaR*^*G4*^ and *PanN*^*G4*^ yielded comparable results, indicating that the *SIFaR*^*G4*^ gene trap captures the functional expression domain of *SIFaR*. These findings indicate that *SIFaR* expression during PSINA is sufficient for wildtype activity cycles and that active phase duration can be regulated by *SIFaR* levels.

### SIFaR in the central brain is necessary for cycles

We had previously reported that electrically silencing only the central brain (CB) subset of *Trpy+* neurons also resulted in loss of brainwide PSINA^28^. This finding had led to the working model of the PSINA engine: a CB-based network of defined neural circuits that generates the patterned activity and relays it across the brain. To ask whether SIFa-SIFaR signaling is consistent with this model, we used a similar approach^28^ to spatially fractionate the broad SIFaR expression domain, which, when silenced in full, leads to complete loss of PSINA (Fig.2h) This approach takes advantage of how the fly optic lobes (OLs) are positioned as satellite neuropils, whose physical separation from the CB make it possible to assess distinct spatial participation in PSINA (Fig.1a,d). We restricted GAL4 expression to either the CB or OLs using conditional GAL80 constructs and a mostly OL-specific FLP recombinase^43^. Within this framework (Fig.3a), we drove temporally-targeted KiR2.1 expression using *PanN*^*G4*^, *Trpγ*^G4^, or *SIFaR*^*G4*^ (Fig.3b-e). Silencing the CB fraction of these expression domains virtually abolished PSINA in both the CB and the OLs (Fig.3c,d). Silencing the OL fraction reduced signal in the OLs by 50-75% and in the CB by ∼30% without altering PSINA cycles in either spatial domain (Fig.3c,e). The non-local effect of OL silencing on signal in the CB may be due to the incompleteness of our spatial fractionation approach or to feedforward/centripetal connections between the OL and the CB^44^. These results indicate that the CB subset of all three expression domains, including *SIFaR*^*G4*^, contain neuronal populations whose activity is necessary for PSINA throughout the brain.

**Figure 3.**
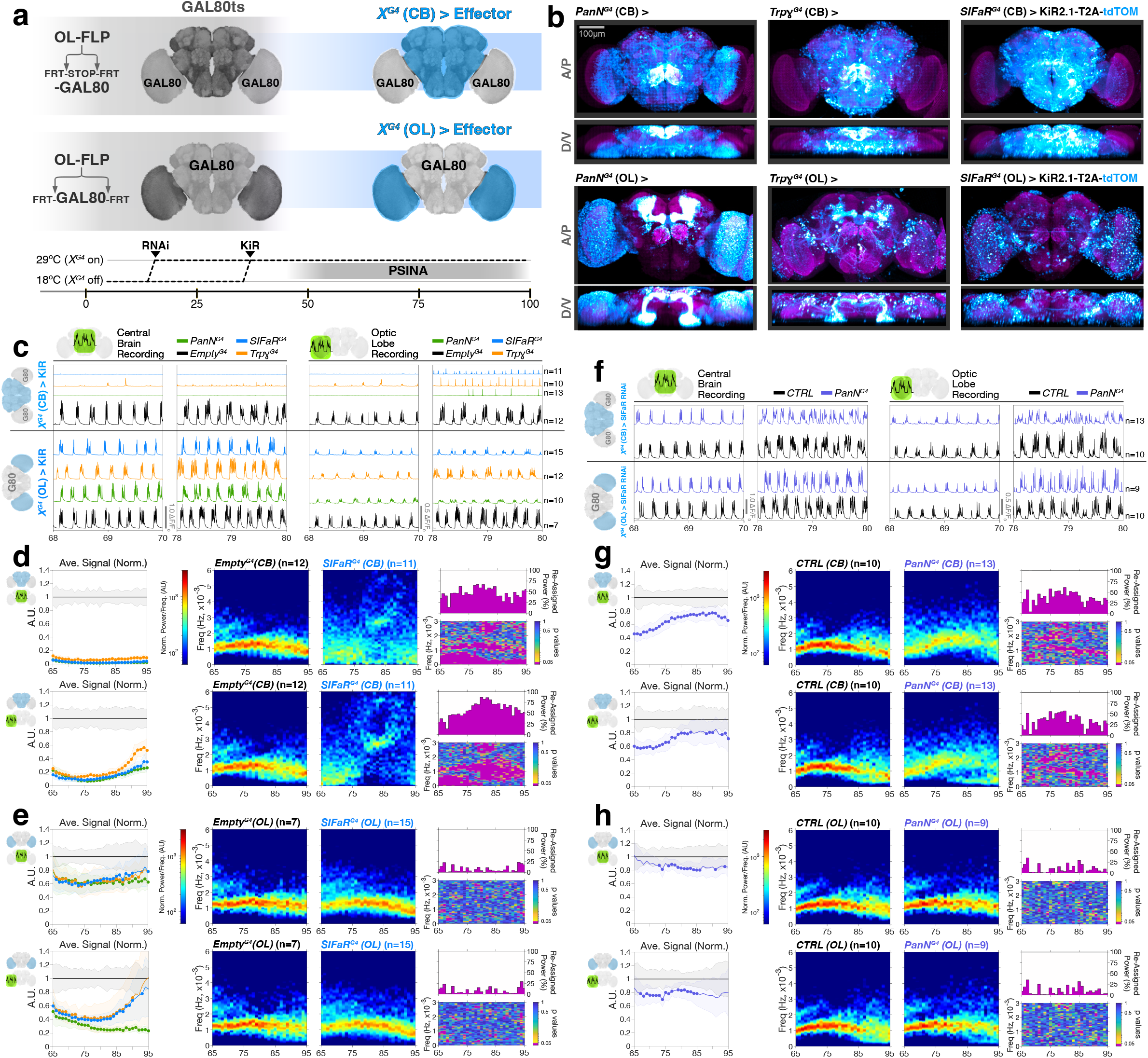
SIFaR in the central brain is necessary for wildtype PSINA cycles. **a**. Spatial fractionation of GAL4 expression. Both schemes carry two variants of *tubP-GAL80*: *GAL80ts* and one of two FLP-responsive conditional alleles. In the optic lobes, OL-FLP (29C07-FLP^43^) either turns on *GAL80* expression by removing the interruption cassette (‘-STOP-’, top) or turns it off by locally excising the FRT-flanked ORF (bottom). Animals are reared at 18^°^C and shifted 29^°^C at in the early pupa to unmask these differential *GAL80* expression domains ahead of PSINA onset; *GAL4*-driven effector expression is disinhibited in the complementary domains (blue). CB, central brain. OL, optic lobe. **b**. Anterior-posterior (A/P) and dorso-ventral (D/V) MIPs of 60hAPF brains with *PanN*-, *Trpγ*-, or *SIFaR*-GAL4 driving *UAS-KiR2.1-T2A-tdTOM* expression principally in the CB (top row) or the OLs (bottom row). *PanN* labeling reveals some posterior OL neurons in the CB-designated fraction and Kenyon cells in the OL-designated fraction. KiR2.1 expression domain detected through tdTOM native fluorescence (cyan-hot LUT); reference stain is N-Cadherin (magenta). Scale bar, 100μm. **c**,**f**. Representative PSINA traces from animals in which KiR2.1 (c) or SIFaR-RNAi (f) expression was targeted to the CB (top set) or OL (bottom set) with the indicated *GAL4*s. CB (left columns) or OL (right columns) recordings are from the same animal in each condition. **d**,**g**. *Left*: Normalized average signal from animals in which KiR2.1 (d) or SIFaR-RNAi (g) expression was spatially targeted to the CB and recorded from the CB (top) or OL (bottom). Genotypes color-matched to (c) and (f); sample sizes indicated in *Right*. Closed circles mark where experimental distributions differ from control (p<0.05) by two-sample KS test. *Right*: Average PSDs from control and experimental conditions described for *Left*; presented as in Fig.1g. **e**,**h**. Data from animals in which KiR2.1 (e) or SIFaR-RNAi (h) expression was spatially targeted to the OLs; presentation as in (d,g).

To assess whether there is also a spatial specialization to gene function, we used the same approach to knock down *SIFaR* in either the CB or the OLs (Fig.3f-h). Consistent with the silencing results, knockdown in either spatial domain reduced PSINA signal. However, only CB knockdown altered cycle across the brain (Fig.3f,g). We conclude that while *SIFaR* is used throughout the brain to support wildtype signal amplitude, it is necessary only in the CB for PSINA cycles.

Taken together, these results are consistent with the CB-based PSINA engine model and also suggest a new layer of amplitude regulation through the brainwide influence of neuropeptide signaling.

### SIFa neuron activity patterns brainwide PSINA cycles

We sought to determine the relationship between the activity of SIFa neurons and brainwide PSINA. The proximity of the oversized SIFa neuronal soma to the dorsal surface of the head made it possible to record from these cells in the widefield assay using GCaMP6s; we followed brainwide PSINA in parallel with RCaMP1b (Fig.4a, Supplementary Video 2). At the level of cycles, SIFa neuron activity is very similar to PSINA (Fig.4b,c). Closer inspection reveals that SIFa neuron active phases both initiate before and end after PSINA active phases (Fig.4b,d). We compared this SIFa neuron lead and lag relative to PSINA in the CB to an internal control—the lead and lag spread between PSINA in the CB and the OL (Fig.4d). Whereas the PSINA active phases across the brain were aligned to within a few seconds, SIFa neurons started out with a lead of tens of seconds and a lag of ∼4 minutes relative to PSINA. These spreads decreased steadily until ∼80hAPF, and SIFa neuron active phases continued to envelope those of PSINA until the end of pupal development. The consistent temporal lead of SIFa neuron active phases suggests a direct role for these neurons in patterning brainwide PSINA cycles.

**Figure 4.**
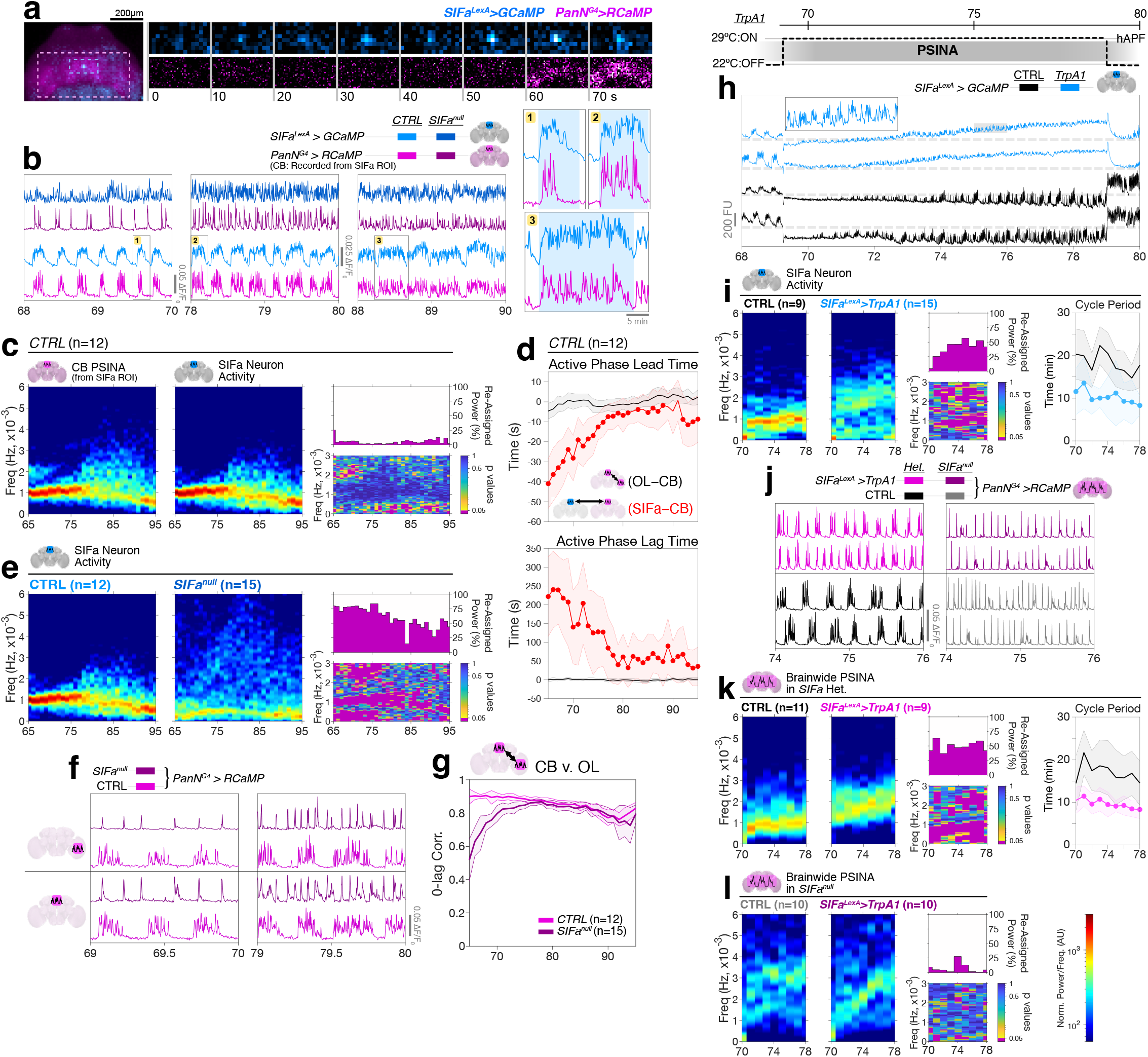
SIFa neuron activity patterns brainwide PSINA cycles. **a-g**. PSINA v. SIFa neuron activity in wildtype and *SIFa* mutant animals. **a**. 2-channel recording of SIFa neuron activity (cyan) and brainwide PSINA (magenta). *Left:* Pan-neuronal>RCaMP and SIFa neuron>GCaMP overlay of pupal head at 69.5 hAPF. *Right:* Progressive frames from boxed span #1 in (b). Color-matched ROIs from *left* shown; intensities adjusted to display fluorescence changes. **b**. *Left:* Representative traces for the indicated conditions. PSINA, as reported by pan-neuronal RCaMP, was recorded from the SIFa neuron ROI in the central brain (CB). *Right:* Expanded views for numbered boxes in *left*; SIFa neuron active phases marked in light blue. **c**. PSD comparison of CB PSINA, as reported by pan-neuronal RCaMP, and SIFa neuron activity in control animals. **d**. Active phase lead (*top)* and lag *(bottom)* time differences between left optic lobe (OL) and CB (black) or SIFa neurons and CB (red) in control animals. ∼30 data points per hourly distribution compiled from 12 data sets; shaded areas, s.d. Closed circles mark where the red distributions differ from black (p<0.05) by two-sample KS test. **e**. PSD comparison of SIFa neuron activity in control and *SIFa* mutant animals; sample sizes as indicated. **f**. Representative PSINA traces recorded from the left OL (*top pair)* or the SIFa neuron ROI in the CB *(bottom pair)* in control (magenta) or *SIFa* mutant (purple) animals. **g**. Zero-lag correlation of PSINA recorded from SIFa neuron ROI in the CB and the left OL in control (magenta) and *SIFa* mutant (purple) animals; sample sizes as indicated. **h-l**. TrpA1 activation of SIFa neurons **h**. *Top:* Temperature schedule used. *Bottom:* SIFa neuron activity response to temperature shift, without (black) and with (blue) TrpA1. *Inset:* Expanded view of gray highlighted span. In our hands, both baseline and change in fluorescence recorded from GCaMP6s varies inversely with temperature. When shifted from 22 to 29^°^C, SIFa neurons in control animals move to a lower baseline (compare to dashed lines marking the baseline at 22^°^C), and, following a pause of ∼1hr, resume periodic activity cycles at a lower signal amplitude. TrpA1-expressing SIFa neurons respond to the temperature up-shift with a graded fluorescence increase which builds up until the down-shift back to 22^°^C. **i**. *Left:* PSD comparison of SIFa neuron activity at 29^°^C without and with TrpA1; sample sizes as indicated. *Right:* SIFa neuron cycle period for the same conditions. Shaded areas, s.d. Closed circles mark where experimental distributions (blue) differ from control (black) (p<0.05) by two-sample KS test. **j**. Representative brainwide PSINA traces at 29^°^C for the indicated genotypes. **k**. *Left:* PSD comparison of brainwide PSINA activity at 29^°^C without and with TrpA1; sample sizes as indicated. *Right:* Brainwide PSINA cycle period for the same conditions. Shaded areas, s.d. Closed circles mark where experimental distributions (magenta) differ from control (black) (p<0.05) by two-sample KS test. **l**. PSD comparison of brainwide PSINA activity at 29^°^C without and with TrpA1 in *SIFa*^*null*^ animals; sample sizes as indicated.

Using the same imaging approach, we studied how loss of *SIFa* affects SIFa neuron activity. We found that SIFa neurons also lose their activity cycles in *SIFa* mutant animals (Fig.4b,e); active and silent phases are replaced by an apparently unstructured train of sweeps (Fig.4b). Thus, the SIFa neuropeptide regulates the activity patterns of SIFa neurons themselves. The requirement for *SIFaR* expression in SIFa neurons (Fig.2f,g) indicates that this is achieved, at least in part, through direct autocrine signaling; below, we explore whether SIFa neurons receive feedback from other neurons.

We next asked whether loss of *SIFa* has differential effects on PSINA across different parts of the brain. Comparing PSINA recorded from the CB and the OL in controls and *SIFa* mutants, we found that the activity remains highly correlated across these different regions in both genotypes (Fig.4f,g). The persistence of the global coordination of PSINA in the absence of *SIFa* suggests that SIFa-SIFaR signaling operates upstream of an independent brainwide activity template^28^.

We studied the patterning roles of SIFa neurons and SIFa-SIFaR signaling in two gain-of-function approaches. First, we overexpressed SIFaR with *SIFaR*^*G4*^ (Extended Data Fig.6). This increased the active phase duration of both SIFa neuron activity and brainwide PSINA by >2-fold (Extended Data Fig.6b,c). SIFa neuron active phases still enveloped those of PSINA with SIFaR overexpression (Extended Data Fig.6a,d). We did observe some hourly changes in the lead and lag times with overexpression (Extended Data Fig.6d); however, these changes were not robust enough to establish whether SIFaR expression levels regulate relative timing of active phases. Taken together with the increase in *SIFaR*^*G4*^ expression during pupal development (Extended Data Fig.4), these results suggest that, in the wildtype, SIFaR levels control the increase in the active phase duration over the course of PSINA.

As a second gain-of-function approach, we used the thermogenic cation channel TrpA1^45^ to activate SIFa neurons (Fig.4h-l). TrpA1-expressing SIFa neurons responded to the activating temperature step with a gradual fluorescence increase; low amplitude activity cycles were visible on top of this graded response (Fig.4h:*Inset*). Compared to controls, the cycle frequency of activated SIFa neurons increased by ∼80% (Fig.4i), resulting in a corresponding reduction in the period of ∼40%. The lower amplitude of TrpA1-activated SIFa neuron cycles precluded reliable two-channel imaging, so we recorded the response of brainwide PSINA in separate experiments (Fig.4j-l). We found that PSINA cycles were similarly sped up in response to SIFa neuron activation (Fig.4j,k). We did not observe any significant changes to brainwide PSINA when SIFa neurons were activated in *SIFa* mutants (Fig.4j,l). These results indicate that SIFa neurons can set the cycle period of both their own activity and that of brainwide PSINA in a SIFa-dependent manner.

The only other gene we know to have a specific effect on the cycle period is the phospholipase Cβ (PLCβ) *NorpA*^25^, an important downstream effector of GPCR signaling: The period in *norpA* nulls is significantly longer, ranging from 200 to 125% of controls between 65-90hAPF (Extended Data Fig.7). Expression of *UAS-NorpA* in SIFa neurons during pupal development fully restores the cycle period in *norpA* nulls (Extended Data Fig.7), indicating that this intracellular signaling factor is important for the patterning role of SIFa neurons.

Based on this set of results, we conclude that SIFa neuron activity, which itself is shaped through the interactions of SIFa, SIFaR, and NorpA, patterns PSINA cycles by modulating the brainwide activity template.

### SIFa neuron activity is regulated by *Trpγ+* neurons

Next, we studied how SIFa neurons interact with the *Trpγ+* population. During pupal development, *Trpγ* is expressed in ∼2,000 neurons, including the SIFa set^28^. Electrically silencing *Trpγ+* neurons in the CB eliminates brainwide PSINA (Fig.3 and ^28^). To ask whether silencing non-SIFa *Trpγ+*, or *Trpγ±* neurons, alters SIFa neuron activity, we devised a set difference approach (Fig.5a,b): Using the GAL4-UAS system, we expressed KiR2.1 in *Trpγ+* neurons while excluding SIFa neurons from silencing with LexA-LexAop driven GAL80. The published, cell-type-specific *SIFa*^*LexA*^ driver^35^ (used for SIFa neuron recordings in Fig.4) uses the transcriptional activation domain of GAL4 and was therefore not compatible with our approach. As an alternative, we identified two other drivers, *SIFa*^*LexA.2*^ and *SIFa*^*LexA.3*^, which make use of the GAL80-insensitive p65 activation domain and are expressed in SIFa neurons during pupal development (Extended Data Fig.8a,b). Using these drivers in combination to drive GAL80 and GCaMP6s (Fig.5b, Extended Data Fig.8c), we were able to both suppress KiR2.1 expression in SIFa neurons and record their activity during PSINA (Fig.5c-e). Silencing *Trpγ±* neurons had two major effects on SIFa neuron activity: First, the silent phases are not as clearly defined as they are in controls: Instead of returning to baseline; SIFa neurons continue to engage in sweeps between major active phases (Fig.5c). This altered pattern, which manifests as broadening of the characteristic frequency band toward higher values in the PSD representation (Fig.5e), is consistent with the loss of an inhibitory input from *Trpγ±* neurons due to KiR2.1 silencing. Secondly, past ∼80hAPF, active phase duration does not increase as dramatically as in controls (Fig.5d), leading to a reduction in both duty cycle and average signal beyond this time point. This suggests that the SIFaR-dependent active phase extension (Fig.6) is acting through *Trpγ±* neurons. In sum, we conclude that *Trpγ±* neurons regulate SIFa neuron activity.

**Figure 5.**
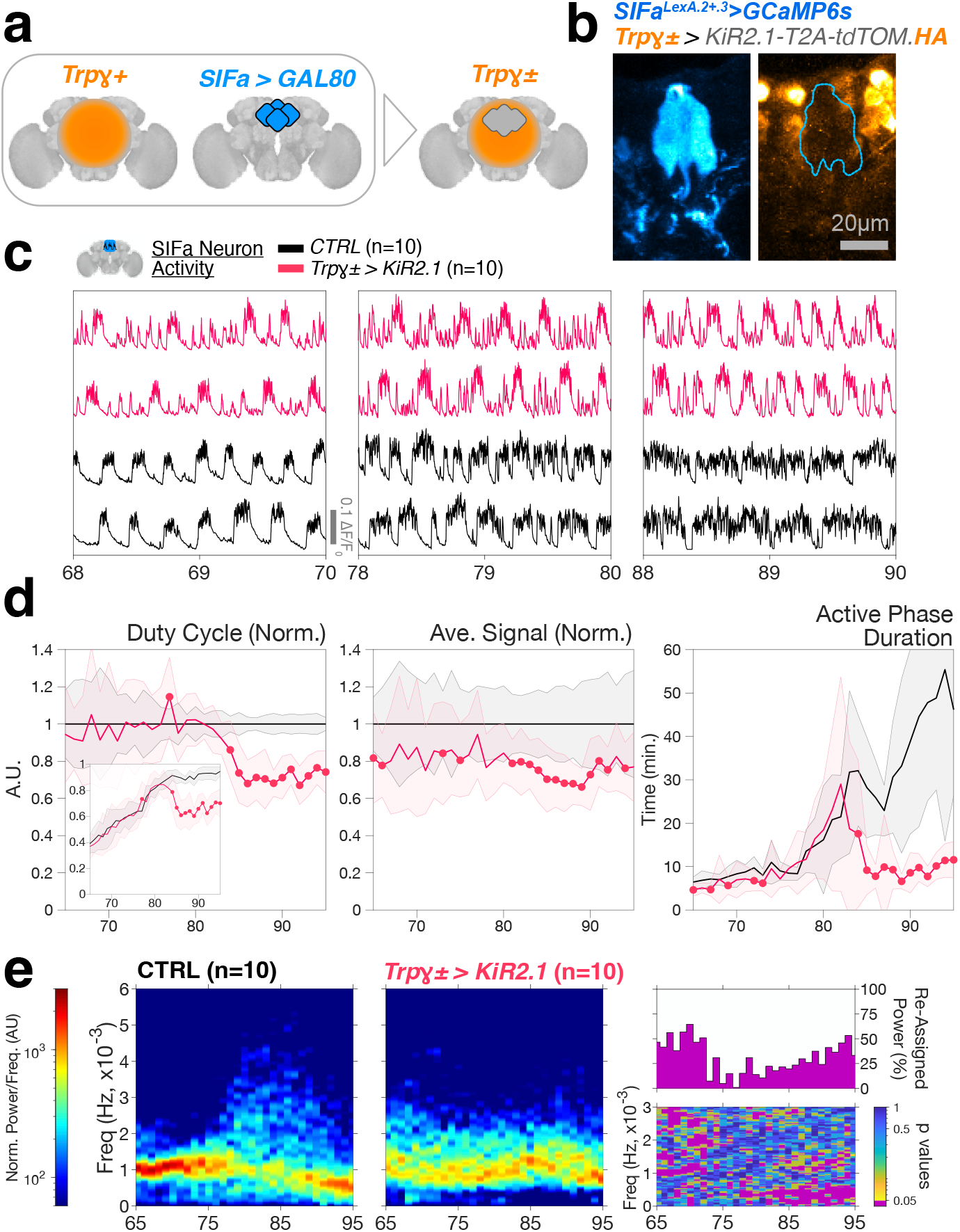
SIFa neuron activity is regulated by *Trpγ+* neurons. **a**. Set Difference: The composite *Trpγ±* driver is generated by combining *Trpγ*^*G4*^ with *SIFa*^*LexA*^ > *LexAop-GAL80*. **b**. MIP of the PI region from a 72hAPF brain with *SIFa*^*LexA.2+.3*^ driving *LexAop-GCaMP6s* (anti-GFP, cyan-hot LUT) and *Trpγ±* driving *UAS-KiR2.1-T2A-tdTOM.HA* (anti-HA, orange-hot LUT); cyan dashed outline marks SIFa soma location in right panel. See Extended Fig.8c for full brain MIP. **c**. Representative traces of SIFa neuron activity in CTRL animals (black) or with *Trpγ±* driving *UAS-KiR2.1* (red). **d**. Normalized and true-valued (inset) duty cycle, normalized average signal, and true-valued active phase duration for the conditions and sample sized indicated in (c). Shaded areas, s.d. Closed circles mark where the experimental distributions differ from CTRL (p<0.05) by two-sample KS test. **e**. Average PSDs from control and experimental animals; presented as in Fig.1g.

**Figure 6.**
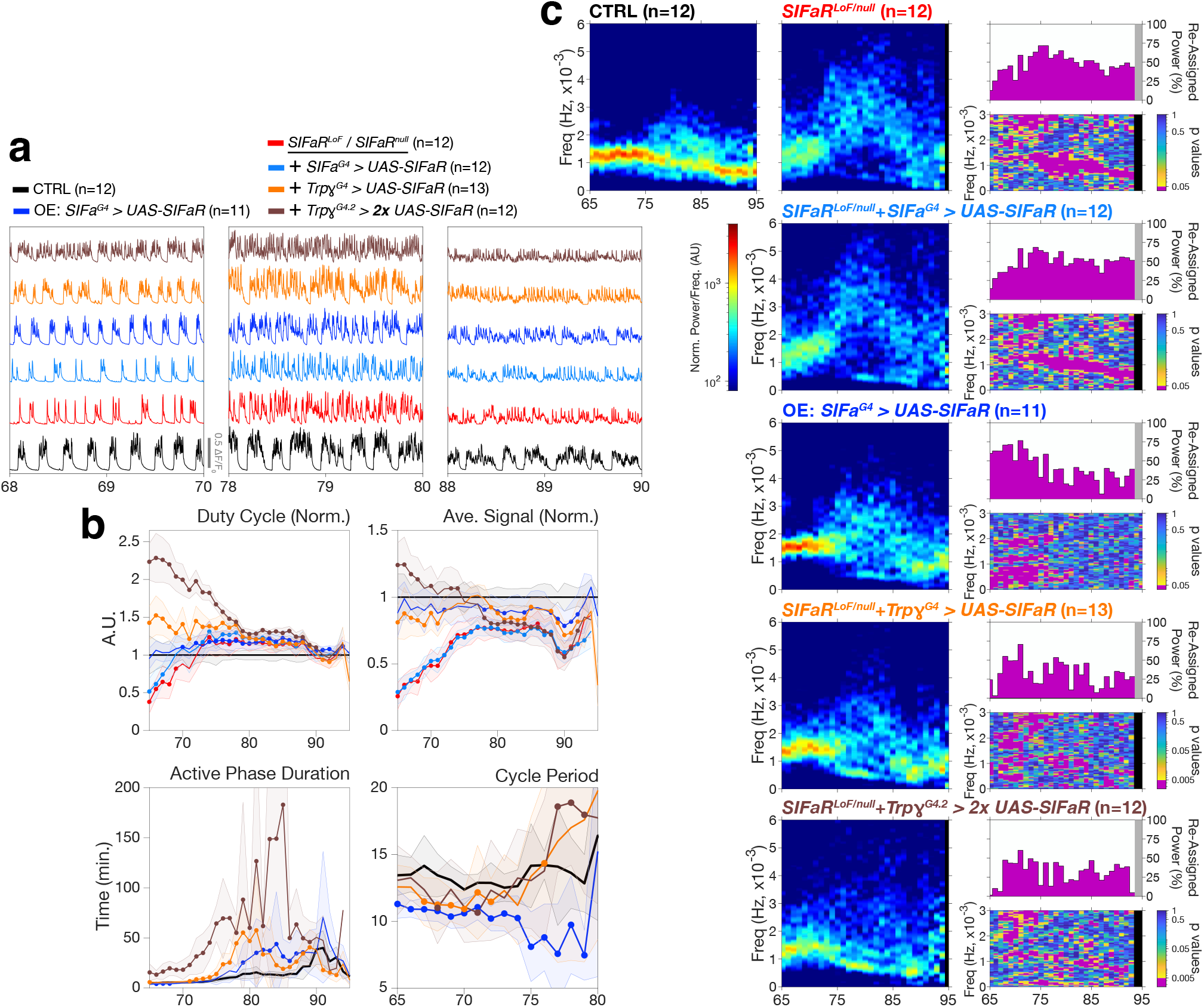
*SIFaR* expression in *Trpγ+* neurons is sufficient to restore brainwide PSINA cycles. **a**. Representative brainwide PSINA traces for the indicated conditions. ‘OE’; overexpression. **b**. Normalized duty cycle and average signal, true-valued active phase duration and cycle period for the conditions and sample sized indicated in (a). Shaded areas, s.d. Closed circles mark where the experimental distributions differ from CTRL (p<0.05) by two-sample KS test. **c**. Average PSDs from control and experimental animals; presented as in Fig.1g.

### SIFaR expression in *Trpy*+ neurons is sufficient to rescue PSINA cycles

The results presented thus far regarding where SIFaR expression is necessary (i.e., *PanN*-, *SIFaR*-, *Trpγ-*, and *SIFa*-GAL4 expression domains., Fig.2f,g) and sufficient (i.e., *PanN*- and *SIFaR*-GAL4 expression domains; Extended Data Fig.5) are consistent with the possibility that *SIFaR* expression in just the four SIFa neurons is sufficient for wildtype PSINA patterning. We found that this is not the case; *SIFa*^*G4*^ driven *UAS-SIFaR* did not restore wildtype PSINA in *SIFaR* mutants (Fig.6). We confirmed the efficacy of this cell-type-specific driver in an overexpression experiment (Fig.6:*’OE’*): Between 65-80hAPF, the cycle frequency increased by ∼37% (Fig.6c), resulting in a corresponding decrease in the period of ∼25% (Fig.6b). Later, from 80-90hAPF, cycle structure was more variable, as seen in the broader frequency spread in the PSD representation (Fig.6c), and the average active phase duration was longer (Fig.6b). These effects—increased cycle frequency and extended active phases—are well-aligned with results from SIFa neuron activation (Fig.4h-l) and *SIFaR*^*G4*^*-*driven overexpression (Extended Data Fig.6), respectively, and are consistent with autonomous SIFaR expression levels regulating the activity patterns of SIFa neurons.

Given that wildtype SIFa neuron activity requires input from *Trpγ±* neurons (Fig.5), we next asked if *SIFaR* expression in the *Trpγ+* population is sufficient to rescue PSINA in *SIFaR* mutants. We found that *Trpγ*^*G4*^ driven *UAS-SIFaR* restored activity cycles (Fig.6a). Compared to controls, PSINA in these animals started out with a lower cycle period and maintained higher duty cycle and longer active phases for most of the duration of observation. These differences were similar to what we had observed with PanN- and *SIFaR*-GAL4 rescue, which had to be titrated down to match wildtype PSINA (Extended Data Fig.5). Consistent with this overexpression inference, increasing the dosage of *Trpγ*^*G4*^ driven *UAS-SIFaR* (Fig.6:*’2x’*) in the context of rescue amplified the differences in duty cycle and active phase duration (Fig.6b). We conclude that *SIFaR* expression in the full *Trpγ+* domain, i.e., SIFa and *Trpγ±* neurons, is sufficient to pattern the PSINA activity cycles.

### SIFa neurons pattern PSINA by modulating *Trpy*+ neurons

Previously, we had shown that brainwide PSINA mirrors the activity recorded from *Trpγ+* neurons^28^ in *trpγ* mutants. As the *Trpγ+* neuron activity reflects the cell autonomous phenotype of the *trpγ* mutation, this result indicated that the *Trpγ+* population acts as an activity template for the rest of the brain. The findings presented here predicted that disruption of *Trpγ+* neuron activity underlies the changes to brainwide PSINA seen in *SIFa* mutants. This is what we observed using two-channel imaging: In *SIFa* mutants, *Trpγ+* neurons exhibit the same activity phenotypes we described for brainwide PSINA: the two signals are nearly identical (Fig.7a-c). We conclude that SIFa neurons pattern the cycles of PSINA by modulating the activity of *Trpγ±* neurons through SIFa-SIFaR signaling (Fig.7d).

**Figure 7.**
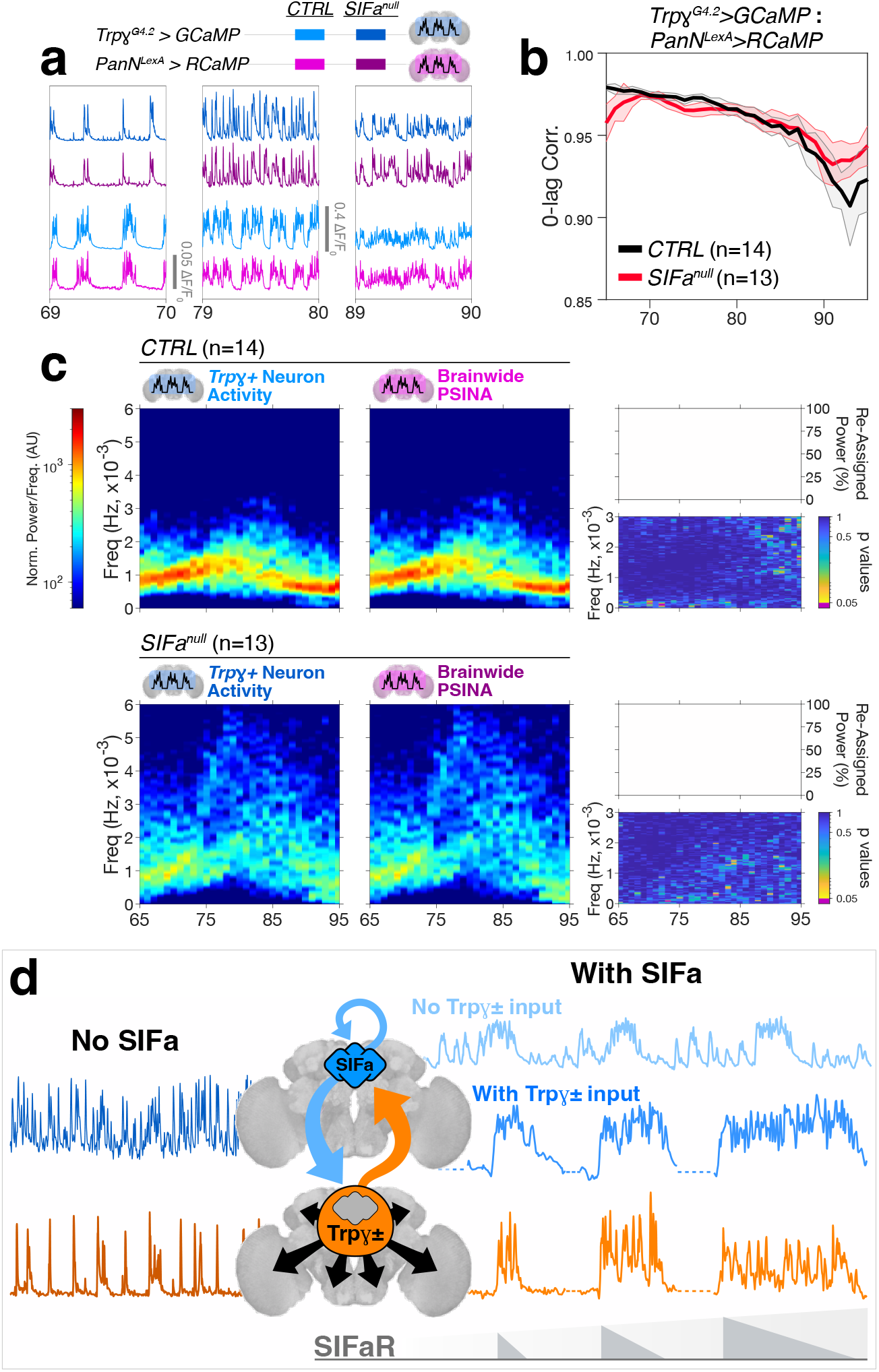
SIFa neurons pattern PSINA by modulating *Trpγ+* neurons. **a-c**. PSINA v. *Trpγ+* neuron activity in wildtype and *SIFa* mutant animals. **a**. Representative traces from 2-channel recordings of *Trpγ+* neuron activity and brainwide PSINA in control and *SIFa* mutant animals. **b**. Zero-lag correlation of *Trpγ+* neuron activity to brainwide PSINA in control and *SIFa* mutant animals. **c**. PSD comparison of *Trpγ+* neuron activity and PSINA in control (top) and *SIFa* mutant (bottom) animals; sample sizes as indicated. **d**. Model of the core PSINA engine. In the absence SIFa signaling (left, ‘No SIFa’), neither SIFa (blue) nor *Trpγ±* (magenta) neurons have stereotyped cycles. SIFa-SIFaR signaling patterns both activity profiles into cycles (right, ‘With SIFa’). SIFa neurons regulate their activity through autocrine signaling, and they receive additional input from *Trpγ±* neurons. Brainwide PSINA follows the *Trpγ±* neuron activity template. SIFaR expression (light gray gradient ramp) and activity levels (dark gray triangles) control the duration of active phases. See *Discussion* for more details.

## DISCUSSION

We report that the neuropeptide gene *SIFa* and its receptor *SIFaR* are required in the developing central brain to establish the characteristic activity cycles of PSINA (Fig.1-3). SIFa neurons themselves have cycles which initiate before brainwide activity (Fig.4b,d). Consistent with this implied lead role in patterning PSINA, activating SIFa neurons increases the frequency of brainwide cycles (Fig.4). Additionally, restoring NorpA expression in just these four cells rescues the reduction in cycle frequency seen in *norpA* mutants (Extended Data Fig.7). *SIFaR* is necessary in SIFa neurons for wildtype brainwide PSINA (Fig.2f,g and Extended Data Fig.3e) and *SIFa* is autonomously required for SIFa neuron activity (Fig.4b,e), indicating that these neurons are autocrine regulated. Wildtype SIFa neuron activity requires additional input from *Trpγ±* neurons (Fig.5). In turn, SIFa neurons pattern the cycles of brainwide PSINA by modulating the activity of *Trpγ+* neurons (Fig.6,7).

These results build on previous work^28^ and suggest a working model of how the core *PSINA engine*—a set of genetically-defined neural circuits in the central brain—patterns brainwide developmental activity (Fig.7d): SIFa-SIFaR signaling transforms the autonomous oscillations of SIFa neurons into cycles. In electrical isolation, the silent phases of SIFa neuron cycles are interrupted by sweeps and the active phases do not extend appreciably over the course of development. SIFa-SIFaR signaling also patterns *Trpγ±* neuron oscillations into cycles. *Trpγ±* neuron active phases grow progressively longer, in parallel with the developmental increase in SIFaR expression (Extended Data Fig.4), suggesting that receptor availability and activation-dependent desensitization regulate this feature of PSINA. Next, *Trpγ±* neurons feedback onto SIFa neurons to establish true silent phases and extend their active phases. Finally, brainwide PSINA follows the *Trpγ±* neuron activity template.

The SIFa-SIFaR dependent patterning of SIFa neuron activity has a precedent in one of the best characterized central pattern generating (CPG) circuits—the crustacean stomatogastric ganglion (STG). In the crab, *Cancer borealis*, SIFa re-organizes the rhythms of the two interdependent CPGs driving food intake and processing^46^. Specifically, one defined neuronal type within these circuits responds to SIFa by adding a low frequency bursting pattern on top of an existing high frequency rhythm^47^. In the STG, which is subject to significant neuromodulation^48^, the response to SIFa depends on both the concentration of the neuropeptide as well as the physiology of the responding cells and circuits^49^. Our observations are consistent with this expected diversity of responses: In the optic lobes, which appear to follow the central brain generated activity template (Fig.3 and ^28^), SIFaR acts as a signal amplifier (Fig.3h). By contrast, in SIFa and *Trpγ+* neurons, both of which may be intrinsically active populations, SIFaR patterns activity into cycles.

The molecular mechanisms of SIFa-SIFaR neuromodulation are not known. Our results implicate the PLCβ *NorpA*^25^ (Extended Data Fig.7) as a candidate effector of SIFaR signaling. Most neuropeptide receptors, including SIFaR, are GPCRs. The primary transducer of GPCR signaling is the heterotrimeric G protein complex comprising a Gα subunit and a Gβγ dimer^50^. In response to receptor activation, Gα dissociates from Gβγ to interact with downstream signaling effectors, which, for the Gαq class of Gα proteins, are PLCβs. Recruitment of PLCβ to the plasma membrane ultimately leads to the activation of protein kinase C and mobilization of Ca^2+^ from intracellular stores. NorpA may be the principal effector of SIFaR signaling; however, PSINA cycles remain intact in *NorpA* nulls (Extended Data Fig.7). There is a second PLCβ in the fly, Plc21C, which can act redundantly with NorpA^51^. It will be possible to resolve this matter and further develop SIFa neurons into an *in vivo* model for neuropeptide signaling using the experimental access established in this work.

Neuropeptides are one of the largest and possibly the oldest class of neuro-active compounds^52,53^. Bilaterian genomes contain many dozens to over 100 neuropeptide genes; this diversity, along with a similar complement of GPCRs, underscores the neurophysiological potential of the chemical connectome^54–56^. In the developing brain, neuropeptides have been implicated in neurogenesis, morphogenesis, migration, survival, and circuit assembly^57–59^. While predicted^60^, a direct involvement of neuropeptide signaling in developmental activity had not been demonstrated until the present study.

A defining feature of neuropeptide signaling is extra-synaptic volume transmission. This mode of communication frees neuropeptidergic cells from the constraints of specific post-synaptic contacts, giving them the reach and flexibility to integrate internal and external stimuli to regulate virtually every physiological and behavioral process in the adult^52,61–63^. During development, the same synapse-free access to neuromodulation would allow neuropeptide signaling to persist through on-going synaptogenesis, resolving the conceptual tension borne by developmental activity as a process tasked with refining synaptic connections it depends on. The possible precedence of neuropeptide signaling in developmental activity also aligns, in ontogeny recapitulates phylogeny fashion, with the chemical brain hypothesis^64^. According to this view, neuropeptide signaling, which may predate metazoan origins^65^, organized the earliest neural networks prior to the emergence of synaptic transmission, after which neuropeptides took on the more familiar modulatory role.

Activity in the developing mammalian brain has been studied in discrete regions, and, in some cases, circuits or mechanisms which serve as local pattern generators have been identified^1^. How these distinct, local activity profiles are coordinated to mediate circuit integration^66^ across disparate areas of the brain is not known. In the much smaller fly system, we show that cross brain coordination is driven by a CPG and relay circuit, the PSINA engine, which minimally comprises the SIFa and *Trpγ±* neurons. Notably, silencing the central PSINA engine reveals local oscillators in the optic lobes (Fig.3c), suggesting that even at the scale of the fly, global coordination of local foci of activity is a built-in feature. SIFa neurons reside in the PI, the neuropeptide and hormone center of the fly brain, and functional homolog of the mammalian hypothalamus. It was recently reported that hypothalamic peptidergic neurons project into nearly all areas of the mouse brain^23^, confirming that the mammalian nervous system has the infrastructure to mediate long-range neuropeptide-mediated coordination; it remains to be seen whether this infrastructure is used during development.

Altering developmental activity is expected to affect behavior, and changes to spontaneous network activity in the developing mammalian cortex are thought to be linked to neurodevelopmental disorders^2^. In the fly, disruption of SIFa-SIFaR signaling affects a range of behaviors; whether these phenotypes are due to loss of neuropeptide signaling in the adult or during development is a matter of interest^67^. The advantages of the fly and our improving handle on the PSINA engine will make it possible to address how PSINA contributes to the emergence neurotypical behavior.

## ACKNOWLEDGEMENTS

We thank J. M. Donlea, M. A. Frye, E. Marcus, P. Sanfilippo, S. L. Zipursky, and members of the Akin Lab for discussions, support and feedback on the manuscript; J. A. Veenstra for sharing SIFa antibodies; D. Hattori for the KiR2.1 flies; A. Nern and G. M. Rubin for the 29C07-FLP flies; B. Deng and Y. Rao for the SIFa[attP] flies; and C. G. Vecsey for a number of fly lines. OA is supported by NIH grant R01NS123376 and W. M. Keck Foundation Junior Faculty Award.

## METHODS

### Experimental model and subject details

Flies were reared at 18°C, 25°C, or 29°C on standard cornmeal/molasses medium; developmental time was matched to the 25°C standard (1x) using relative rates of 0.5x and 1.25x for 18°C and 29°C^68^, respectively. Pupal development was staged relative to white pre-pupa formation (w.p.p., 0hAPF) or head eversion (h.e., 12hAPF). GAL4/UAS^69^ and LexA/LexAop^70^ expression systems were used to drive cell-type-specific transgene expression. The available molecularly-defined null allele of *SIFaR, SIFaR*^*[attP]*55^, appeared to be homozygous lethal. However, complementation tests indicated that a second site mutation was the cause of the lethality, and we recovered a viable SIFaR^[attP]^ line through out-crosses. Complete genotypes used in each experiment can be found in Supplementary Table 1.

### Widefield imaging

Bottle-scale (∼100 virgins, 10-20 males) crosses were setup for each genotype of each experimental set and reared at 18°C-29°C depending on experimental design. Pupae were genotyped and staged at w.p.p. or h.e. All data shown were collected from female pupae; mutant phenotypes of *SIFa* and *SIFaR* (Fig.1) were also confirmed in males (not shown). Imaging sessions typically started ∼50hAPF and lasted until eclosion (∼100hAPF). To prepare the pupae for imaging, the cuticle around the head was removed with fine forceps and the animals were fixed to a metal plate (McMaster-Carr) with double-sided adhesive tape (3M). The metal plate was placed in a custom environmental chamber to maintain temperature and humidity. This chamber comprised: a PTC1 temperature-controlled breadboard (Thorlabs) to maintain sample temperature at a designated set-point; a set of four 35-mm dishes filled with deionized water to maintain humidity; and a 150-mm petri-dish lid to provide enclosure. To improve imaging quality, a large format coverslip was fitted into a rectangular opening made in the lid. This coverslip sat in the optical path, between the pupae and the objective lens, and was treated with Barbasol shaving cream (Perio) to prevent condensation during temperature shifts.

Images were acquired using either a Zeiss (1- and 2-channel imaging) Axio Zoom.V16 epifluorescence microscope equipped with a Retiga R1 CCD Camera (QImaging) and X-Cite TURBO 6-Channel LED Light Source (Excelitas Technologies), or a Leica (1-channel imaging) M165FC epifluorescence microscope with a Leica.K5 Camera and CoolLED 3-Channel Light Source (CoolLED). Images were acquired at 0.5 Hz with 100ms exposure time for GCaMP6s and 200ms exposure time for jRCaMP1b (5-20mW under objective power). Acquisition was controlled by Slidebook software (Intelligent Imaging Innovations) on the Zeiss microscope and LAS X (Leica Inc) on the Leica microscope. Time series were processed with Fiji^71^(ImageJ) and analyzed using MATLAB (Mathworks); see below.

### Immunofluorescence and confocal microscopy

Brains were dissected in cold Schneider Medium (Gibco 21720–024) and fixed with 4% v/v PFA (Electron Microscopy Sciences 15710) in Schneider Medium for 20 min at room temperature. Brains were then washed out of fixative into PBS (Quality Biological), solubilized in PBST (0.5% Triton-X100 (Sigma T9284) in PBS) for 1 h, and blocked in PBTN (5% Normal Donkey Serum (Jackson ImmunoResearch no. 017-000-121) in PBST) for 1-2 h, all at room temperature. Brains were sequentially incubated in primary and secondary antibodies diluted in PBTN for 24–48 h at 4 °C, with at least 3 washes through PBST over 2 h at room temperature in between and afterwards. Brains were post-fixed with 4% v/v PFA in Schneider Medium for 20 min at room temperature, followed by multiple washes into PBST over 10 min. Brains were finally transferred to Everbrite mounting medium (Biotium 23001) and mounted on to slides for imaging.

Primary antibodies and dilutions used in this study were as follows: chicken anti-GFP (Abcam ab13970, 1:1,000), mouse monoclonal anti-V5 (Novus Biologicals NBP2-52703–0.2 mg, 1:150), rat monoclonal anti-Ncad (DSHB DN-Ex 8-c, 1:100), mouse monoclonal anti-Brp (DSHB nc82-c, 1:150), and rabbit polyclonal anti-SIFamide^30^ (1:1,000).

Secondary antibodies and dilutions used in this study were as follows: Alexa 488 donkey polyclonal anti-chicken (Jackson ImmunoResearch 703-545-155, 1:400), Alexa 488 donkey polyclonal anti-mouse (Jackson ImmunoResearch 715-545-151, 1:400), Alexa 488 donkey polyclonal anti-rabbit (Jackson ImmunoResearch 712-545-153, 1:400), Alexa 568 donkey polyclonal anti-mouse (Jackson ImmunoResearch 715-575-150, 1:400), Alexa 568 donkey polyclonal anti-rabbit (Invitrogen A10042, 1:400), Alexa 647 donkey polyclonal anti-rat (Jackson ImmunoResearch 712-605-153, 1:400), and Alexa 647 donkey polyclonal anti-rabbit (Jackson ImmunoResearch 711-605-152, 1:400) Immunofluorescence images were acquired using a Zeiss LSM 780 confocal microscope with 20×/0.8 air, 40×/1.4 oil immersion or 40×/1.2 glycerol immersion objectives (Zeiss). Images were prepared for publication using Fiji^71^(ImageJ).

### Analysis of widefield imaging data

User-defined ROIs from time series were selected in Fiji^71^ (ImageJ). Custom scripts written in MATLAB (Mathworks) were used in all subsequent analysis. Raw fluorescence traces from per frame ROI pixel averages (F) were fit to a baseline (F_0_) and ΔF/F_0_ (signal) was calculated by subtracting the baseline from the raw signal and dividing by the baseline. Between 75-85hAPF, the raw fluorescence recorded from intact pupae exhibits a stereotyped sigmoidal decrease in intensity correlated with the maturation of the cuticle around the head. By comparing this trend, captured by the baseline fit (F_0_), to staged animals, we found that the inflection point of the curve corresponds to ∼80hAPF. In all analyses, this internal, genotype-independent feature was used to synchronize the developmental ages of the animals. The pace of PSINA (e.g., cycle frequency) is temperature sensitive. To compare data collected at different temperatures, we scaled the time axes of the recordings to match the 25°C standard (1x) using the coefficient T_Coef_=100/(-0.0465*T^3^+4.263*T^2^-133.49*T+1500), where T is the recording temperature in degrees Celsius. The formula is derived from the best fit third order polynomial to the measured temperature-dependent duration of metamorphosis^68^. Active phase limits were defined using a threshold set to 4% of average maximum signal recorded from control genotypes with a minimum floor of 0.5% ΔF/F_0_ for low signal-to-noise data (e.g., SIFa neuron recordings).

PSDs were calculated for each hour of recording using a 2-h sliding window^72^. In preparation for Fast Fourier Transform and PSD estimation, each 2-h segment was filtered to isolate target frequencies (standard 120s Gaussian window to remove sweeps or defined low pass cutoffs as noted in Figure legends), centered on the time axis by subtracting its mean, and multiplied with the Hanning window function to reduce spectral leakage. The PSD for each 2-hr segment was normalized relative to its discrete integral over all frequencies; this step effectively removes the contribution of signal amplitude changes to the PSD in order to focus on frequency changes for subsequent comparisons. Each experimental condition thus yielded a distribution of normalized PSD values (as many as replicate number) at each frequency-time coordinate. Statistical tests between different experimental conditions were carried out between these distributions. PSDs from all replicates of an experimental condition were averaged to generate the average PSD panels shown in the Figures.

All statistical analyses were carried out using the non-parametric two-sample Kolmogorov-Smirnov test; p values less than 0.05 were considered significant.

## REFERENCES

1. Blankenship, A. G. & Feller, M. B. Mechanisms underlying spontaneous patterned activity in developing neural circuits. Nature Reviews Neuroscience 11, 18–29 (2010).

2. Wu, M. W., Kourdougli, N. & Portera-Cailliau, C. Network state transitions during cortical development. Nat. Rev. Neurosci. 25, 535–552 (2024).

3. Martini, F. J., Guillamón-Vivancos, T., Moreno-Juan, V., Valdeolmillos, M. & López-Bendito, G. Spontaneous activity in developing thalamic and cortical sensory networks. Neuron 109, 2519–2534 (2021).

4. Thompson, A., Gribizis, A., Chen, C. & Crair, M. C. Activity-dependent development of visual receptive fields. Current Opinion in Neurobiology 42, 136–143 (2017).

5. Kersbergen, C. J., Babola, T. A., Rock, J. & Bergles, D. E. Developmental spontaneous activity promotes formation of sensory domains, frequency tuning and proper gain in central auditory circuits. Cell Reports 41, 111649 (2022).

6. Antón-Bolaños, N. et al. Prenatal activity from thalamic neurons governs the emergence of functional cortical maps in mice. Science 364, 987–990 (2019).

7. Ge, X. et al. Retinal waves prime visual motion detection by simulating future optic flow. Science 373, (2021).

8. Tiriac, A., Bistrong, K., Pitcher, M. N., Tworig, J. M. & Feller, M. B. The influence of spontaneous and visual activity on the development of direction selectivity maps in mouse retina. Cell Reports 38, 110225 (2022).

9. Willshaw, D. J., Von Der Malsburg, C. & Longuet-Higgins, H. C. How patterned neural connections can be set up by self-organization. Proceedings of the Royal Society of London. Series B. Biological Sciences 194, 431– 445 (1976).

10. Matsumoto, N., Barson, D., Liang, L. & Crair, M. C. Hebbian instruction of axonal connectivity by endogenous correlated spontaneous activity. Science 385, eadh7814 (2024).

11. Crair, M. C. Neuronal activity during development: permissive or instructive? Current Opinion in Neurobiology 9, 88–93 (1999).

12. Sernagor, E. & Hennig, M. H. Retinal Waves. 1–12 (2013) doi:10.1016/B978-0-12-397266-8.00041-7.

13. Nakashima, A. et al. Structured spike series specify gene expression patterns for olfactory circuit formation. Science 365, eaaw5030 (2019).

14. Kersbergen, C. J. & Bergles, D. E. Priming central sound processing circuits through induction of spontaneous activity in the cochlea before hearing onset. Trends in Neurosciences 47, 522–537 (2024).

15. Galli, L. & Maffei, L. Spontaneous impulse activity of rat retinal ganglion cells in prenatal life. Science 242, 90–91 (1988).

16. Meister, M., Wong, R. O., Baylor, D. A. & Shatz, C. J. Synchronous bursts of action potentials in ganglion cells of the developing mammalian retina. Science 252, 939–943 (1991).

17. Ackman, J. B., Burbridge, T. J. & Crair, M. C. Retinal waves coordinate patterned activity throughout the developing visual system. Nature 490, 219–225 (2012).

18. Bansal, A. et al. Mice lacking specific nicotinic acetylcholine receptor subunits exhibit dramatically altered spontaneous activity patterns and reveal a limited role for retinal waves in forming ON and OFF circuits in the inner retina. Journal of Neuroscience 20, 7672–7681 (2000).

19. McLaughlin, T., Torborg, C. L., Feller, M. B. & O’Leary, D. D. M. Retinotopic map refinement requires spontaneous retinal waves during a brief critical period of development. Neuron 40, 1147–1160 (2003).

20. Rossi, F. M. et al. Requirement of the nicotinic acetylcholine receptor β2 subunit for the anatomical and functional development of the visual system. Proceedings of the National Academy of Sciences 98, 6453– 6458 (2001).

21. Murcia-Belmonte, V. et al. A Retino-retinal Projection Guided by Unc5c Emerged in Species with Retinal Waves. Current Biology 29, 1149-1160.e4 (2019).

22. Warwick, R. A. et al. Top-down modulation of the retinal code via histaminergic neurons of the hypothalamus. Science Advances 10, eadk4062 (2024).

23. Jiao, Z. et al. Projectome-based characterization of hypothalamic peptidergic neurons in male mice. Nat Neurosci 1–16 (2025) doi:10.1038/s41593-025-01919-0.

24. Carreira-Rosario, A., York, R. A., Choi, M., Doe, C. Q. & Clandinin, T. R. Mechanosensory input during circuit formation shapes Drosophila motor behavior through patterned spontaneous network activity. Current biology : CB https://doi.org/10.1016/j.cub.2021.08.022 (2021) doi:10.1016/j.cub.2021.08.022.

25. Akin, O., Bajar, B. T., Keles, M. F., Frye, M. A. & Zipursky, S. L. Cell-type-Specific Patterned Stimulus-Independent Neuronal Activity in the Drosophila Visual System during Synapse Formation. Neuron 101, 894-904.e5 (2019).

26. Chen, Y. et al. Cell-type-specific labeling of synapses in vivo through synaptic tagging with recombination. Neuron 81, 280–293 (2014).

27. Muthukumar, A. K., Stork, T. & Freeman, M. R. Activity-dependent regulation of astrocyte GAT levels during synaptogenesis. Nature Neuroscience 17, 1340–1350 (2014).

28. Bajar, B. T. et al. A discrete neuronal population coordinates brain-wide developmental activity. Nature 602, 639–646 (2022).

29. Verleyen, P. et al. SIFamide is a highly conserved neuropeptide: a comparative study in different insect species. Biochemical and Biophysical Research Communications 320, 334–341 (2004).

30. Terhzaz, S., Rosay, P., Goodwin, S. F. & Veenstra, J. A. The neuropeptide SIFamide modulates sexual behavior in Drosophila. Biochemical and Biophysical Research Communications 352, 305–310 (2007).

31. Wolff, T. et al. Cell type-specific driver lines targeting the Drosophila central complex and their use to investigate neuropeptide expression and sleep regulation. eLife 14, (2025).

32. Hartenstein, V. The neuroendocrine system of invertebrates: a developmental and evolutionary perspective. Journal of Endocrinology 190, 555–570 (2006).

33. Namiki, S., Dickinson, M. H., Wong, A. M., Korff, W. & Card, G. M. The functional organization of descending sensory-motor pathways in Drosophila. eLife 7, e10806 (2018).

34. Jørgensen, L. M., Hauser, F., Cazzamali, G., Williamson, M. & Grimmelikhuijzen, C. J. P. Molecular identification of the first SIFamide receptor. Biochemical and Biophysical Research Communications 340, 696– 701 (2006).

35. Martelli, C. et al. SIFamide Translates Hunger Signals into Appetitive and Feeding Behavior in Drosophila. CellReports 20, 464–478 (2017).

36. Dreyer, A. P. et al. A circadian output center controlling feeding:fasting rhythms in Drosophila. PLOS Genetics 15, e1008478 (2019).

37. Bai, L. et al. A Conserved Circadian Function for the Neurofibromatosis 1 Gene. CellReports 22, 3416–3426 (2018).

38. Park, S., Sonn, J. Y., Oh, Y., Lim, C. & Choe, J. SIFamide and SIFamide receptor defines a novel neuropeptide signaling to promote sleep in Drosophila. Molecules and cells 37, 295–301 (2014).

39. Sellami, A. & Veenstra, J. A. SIFamide acts on fruitless neurons to modulate sexual behavior in Drosophila melanogaster. Peptides 74, 50–56 (2015).

40. Nässel, D. R. Substrates for Neuronal Cotransmission With Neuropeptides and Small Molecule Neurotransmitters in Drosophila. Frontiers in Cellular Neuroscience 12, 83 (2018).

41. Baines, R. A., Uhler, J. P., Thompson, A., Sweeney, S. T. & Bate, M. Altered electrical properties in Drosophila neurons developing without synaptic transmission. Journal of Neuroscience 21, 1523–1531 (2001).

42. Aso, Y. et al. The neuronal architecture of the mushroom body provides a logic for associative learning. eLife 3, e04577 (2014).

43. Nern, A., Pfeiffer, B. D. & Rubin, G. M. Optimized tools for multicolor stochastic labeling reveal diverse stereotyped cell arrangements in the fly visual system. Proceedings of the National Academy of Sciences 112, E2967–E2976 (2015).

44. Nern, A. et al. Connectome-driven neural inventory of a complete visual system. Nature 641, 1225–1237 (2025).

45. Pulver, S. R., Pashkovski, S. L., Hornstein, N. J., Garrity, P. A. & Griffith, L. C. Temporal dynamics of neuronal activation by Channelrhodopsin-2 and TRPA1 determine behavioral output in Drosophila larvae. Journal of Neurophysiology 101, 3075–3088 (2009).

46. Blitz, D. M., Christie, A. E., Cook, A. P., Dickinson, P. S. & Nusbaum, M. P. Similarities and differences in circuit responses to applied Gly1-SIFamide and peptidergic (Gly1-SIFamide) neuron stimulation. Journal of Neurophysiology 121, 950–972 (2019).

47. Fahoum, S.-R.H. & Blitz, D. M. Neuronal Switching between Single- and Dual-Network Activity via Modulation of Intrinsic Membrane Properties. J. Neurosci. 41, 7848–7863 (2021).

48. Marder, E. Neuromodulation of Neuronal Circuits: Back to the Future. Neuron 76, 1–11 (2012).

49. Blitz, D. M., Christie, A. E., Cook, A. P., Dickinson, P. S. & Nusbaum, M. P. Similarities and differences in circuit responses to applied Gly1-SIFamide and peptidergic (Gly1-SIFamide) neuron stimulation. Journal of Neurophysiology 121, 950–972 (2019).

50. Hanlon, C. D. & Andrew, D. J. Outside-in signaling – a brief review of GPCR signaling with a focus on the Drosophila GPCR family. Journal of Cell Science 128, 3533–3542 (2015).

51. Shieh, B.-H., Sun, W. & Ferng, D. A conventional PKC critical for both the light-dependent and the light-independent regulation of the actin cytoskeleton in Drosophila photoreceptors. Journal of Biological Chemistry 299, (2023).

52. Beets, I. & Watteyne, J. Mapping and decoding neuropeptide signaling networks in nervous system function. Current Opinion in Neurobiology 92, 103027 (2025).

53. Hauser, F., Koch, T. L. & Grimmelikhuijzen, C. J. P. Review: The evolution of peptidergic signaling in Cnidaria and Placozoa, including a comparison with Bilateria. Front. Endocrinol. 13, (2022).

54. Ripoll-Sánchez, L. et al. The neuropeptidergic connectome of C. elegans. Neuron 111, 3570-3589.e5 (2023).

55. Deng, B. et al. Chemoconnectomics: Mapping Chemical Transmission in Drosophila. Neuron 101, 876-893.e4 (2019).

56. Smith, S. J. et al. Single-cell transcriptomic evidence for dense intracortical neuropeptide networks. eLife 8, e47889 (2019).

57. Bakos, J., Zatkova, M., Bacova, Z. & Ostatnikova, D. The Role of Hypothalamic Neuropeptides in Neurogenesis and Neuritogenesis. Neural Plasticity 2016, 3276383 (2016).

58. Li, Z.-H., Li, B.Zhang, X.-Y. & Zhu, J.-N. Neuropeptides and Their Roles in the Cerebellum. International Journal of Molecular Sciences 25, 2332 (2024).

59. Hevesi, Z. et al. Transient expression of the neuropeptide galanin modulates peripheral-to-central connectivity in the somatosensory thalamus during whisker development in mice. Nat Commun 15, 2762 (2024).

60. Hevesi, Z., Hökfelt, T. & Harkany, T. Neuropeptides: The Evergreen Jack-of-All-Trades in Neuronal Circuit Development and Regulation. BioEssays 47, e202400238 (2025).

61. Jékely, G. & Yuste, R. Nonsynaptic encoding of behavior by neuropeptides. Current Opinion in Behavioral Sciences 60, 101456 (2024).

62. Nässel, D. R. & Zandawala, M. Recent advances in neuropeptide signaling in Drosophila, from genes to physiology and behavior. Progress in Neurobiology 179, 101607 (2019).

63. Nässel, D. R. & Zandawala, M. Endocrine cybernetics: neuropeptides as molecular switches in behavioural decisions. Open Biology 12, 220174 (2022).

64. Jékely, G. The chemical brain hypothesis for the origin of nervous systems. Philosophical Transactions of the Royal Society B: Biological Sciences 376, 20190761 (2021).

65. Yañez-Guerra, L. A., Thiel, D. & Jékely, G. Premetazoan Origin of Neuropeptide Signaling. Molecular Biology and Evolution 39, msac051 (2022).

66. Guillamón-Vivancos, T. et al. Input-dependent segregation of visual and somatosensory circuits in the mouse superior colliculus. Science 377, 845–850 (2022).

67. Huang, H., Possidente, D. R. & Vecsey, C. G. Optogenetic activation of SIFamide (SIFa) neurons induces a complex sleep-promoting effect in the fruit fly Drosophila melanogaster. Physiology & Behavior 239, 113507 (2021).

68. Bainbridge, S. P. & Bownes, M. Staging the metamorphosis of Drosophila melanogaster. Journal of embryology and experimental morphology 66, 57–80 (1981).

69. Brand, A. H. & Perrimon, N. Targeted gene expression as a means of altering cell fates and generating dominant phenotypes. Development 118, 401–415 (1993).

70. Lai, S.-L. & Lee, T. Genetic mosaic with dual binary transcriptional systems in Drosophila. Nature Neuroscience 9, 703–709 (2006).

71. Schindelin, J. et al. Fiji: an open-source platform for biological-image analysis. Nat Methods 9, 676–682 (2012).

72. Uhlén, P. Spectral Analysis of Calcium Oscillations. Sci. STKE 2004, (2004).

