## Supplementary Text and Figures for "Four neurons pattern brain-wide developmental activity through neuropeptide signaling"

Extended Data Figure Legends
Extended Data Figures 1-8
Supplementary Videos 1-2
Supplementary Table 1

#### EXTENDED DATA FIGURE LEGENDS

**Extended Data Figure 1. Power Spectral Density based analysis of PSINA; related to Fig.1.**
Data displayed in Fig.1g (*+/SIFa$^1$* v. *SIFa$^1$*; framed in light blue) is used to illustrate the steps of our PSD-based analysis.
**a.** For each fly (e.g., Fly#1) of a genotype (*+/SIFa$^1$* v. *SIFa$^1$*), a 2-h long PSINA segment (light gray traces in first column of plots) centered around each hour of recording (e.g., 80hAPF) was processed (black overlaid traces, see Methods) to prepare for Discrete Fourier Transform ('F') and PSD calculation (second column of plots). Concatenating these hourly PSDs (third column of plots) show how the PSINA signal is distributed across different frequencies from 65-95 hAPF. Average hourly PSDs (bottom row of second column; shaded areas, s.d.) were calculated for each genotype and similarly concatenated to generate the PSD representation (bottom row of third column) shown in main figures. This heat map plot only displays the average and not the spread (e.g., s.d.) of the PSD distribution calculated for each genotype. Note that the heat map values for (Normalized Power / Freq.) are plotted in log scale to better display the PSD dynamic range.
**b.** *First column, top plot:* The 80hAPF PSD distributions from *+/SIFa$^1$* (black, n=12) and *SIFa$^1$* (red, n=15) animals in (a) are overlaid; shaded areas, s.d. Bold line segments (magenta) mark where the *SIFa$^1$* distribution differs from *+/SIFa$^1$* ($p<0.05$) by two-sample KS test.
*First column, center plot:* p values calculated at 80hAPF across the displayed frequency range are plotted. Magenta flood line marks where $p<0.05$; these regions correspond to the bold line segments in the top plot.
*Second column, bottom plot:* Hourly p values are concatenated to show when and at what frequency PSINA in the experimental (*SIFa$^1$*) genotype differs from control (*+/SIFa$^1$*).
*First column, bottom plot:* The fraction of normalized signal power in the control PSD at frequencies where the experimental PSD is significantly different is calculated by taking the ratio of the area under the curve at $p<0.05$ regions to the total area. We defined this fraction, or 100x of this fraction, as percent of control signal power re-assigned to different frequencies. This scalar metric offers a salience measure to the statistical test by describing how much of PSINA power is lost to changes in the PSD.
*Second column, top plot:* Hourly re-assigned power values are concatenated across developmental time.

**Extended Data Figure 2. *SIFa* and *SIFaR* genetic analysis; Related to Fig.1.**
**a.** Duty cycle and average signal of indicated genotypes, normalized to control in Fig.1e. Inset in duty cycle plots true values of metric. Shaded areas, s.d. Closed circles mark where *+/SIFa[1] + Dp* (blue) distribution differs from *SIFa[1] + Dp* (green, reproduced from Fig.1f) ($p<0.05$) by two-sample KS test. The similarity of these data suggests that the differences between genomic rescue and control (Fig.1f) is due to positional effects from the relocated genomic duplication.
**b-d.** *SIFaR* mutant analysis with *SIFaR$^{G4}$*; data presented as in Fig.1e-g.
**e-g.** Double mutant analysis with *SIFaR$^{LoF}$*; data presented as in Fig.1e-g. Statistical tests carried out against double mutant (magenta) in (f,g).
**h-j.** Double mutant analysis with *SIFaR$^{TG4}$*; data presented as in Fig.1e-g. Statistical tests carried out against double mutant (magenta) in (i,j).

**Extended Data Figure 3. Related to Fig.2.**
**a.** MIP of 72hAPF central brain with *SIFa*$^{G4}$ driving *myr::SM-V5* (cyan-hot LUT); brain also stained against SIFa (orange-hot LUT) and N-Cadherin (magenta). Green square marks location of SIFa soma. Scale bar, 50µm.
**b.** MIP of 72hAPF brain with *SIFa*$^{G4-2}$ (i.e., *14F05*$^{G4}$) driving *myr::SM-V5* (cyan-hot LUT); brain also stained against SIFa (orange-hot LUT) and N-Cadherin (magenta). Green square marks location of SIFa soma. Scale bar, 100µm.
**c-e.** Duty cycle and average signal from
**c.** *SIFa* knockdown (Fig.2b,c),
**d.** KiR2.1 expression in SIFa neurons (Fig.2d,e),
**e.** *SIFaR* knockdown (Fig.2f,g) experiments.

**Extended Data Figure 4. *SIFaR*$^{TG4}$ developmental expression domain.**
*Left:* MIPs of brains at the indicated ages with *SIFaR*$^{TG4}$ driving *mCherry.NLS* (native fluorescence, cyan-hot LUT); brain also stained against SIFa (yellow-hot LUT) and N-Cadherin (magenta). Green boxes mark location of SIFa soma panels in *Right*. Raw image stacks acquired with 20X objective using consistent image acquisition settings; *mCherry.NLS* and SIFa channel MIPs displayed with same contrast settings. Scale bar, 100µm.
*Right:* MIPs of SIFa soma (yellow) and *mCherry.NLS* (native fluorescence, cyan-hot LUT) from brains in *Left.* Raw image stacks acquired with 40X objective using consistent image acquisition settings; SIFa channel MIPs displayed with same contrast settings. Arrowheads mark SIFa neuron nuclei. Scale bar, 20µm.

**Extended Data Figure 5. Temporally targeted and tuned rescue of PSINA in *SIFaR* mutants.**
**a.** Temperature schedule and genotypes.
**b-d.** *GAL80ts* targeted *GAL4-UAS* rescue of *SIFaR* phenotype at 28$^{o}$C.
**b.** PSINA traces, color-matched to genotypes in (a).
**c.** Duty cycle, average signal, and active phase duration, all normalized to control genotype (black). Plots color-matched to genotypes in (a); sample sizes indicated in (d). Shaded areas, s.d. Closed circles mark where experimental distributions differ from control ($p<0.05$) by two-sample KS test.
**d.** Average PSDs from control and experimental animals; presented as in Fig.1g.
**e-g.** *GAL80ts* targeted *GAL4-UAS* rescue of *SIFaR* phenotype at 24$^{o}$C; presented as in (b-d).

**Extended Data Figure 6. SIFaR overexpression increases active phase duration; Related to Fig.4.**
**a.** *Left:* Representative traces of SIFa neuron activity (cyan and dark blue) and brainwide PSINA (magenta and purple) in CTRL animals or with SIFaR overexpression. PSINA, as reported by *SIFaR*$^{G4}$*>RCaMP*, was recorded from the SIFa neuron ROI. *Right:* Expanded views for numbered boxes in *left*; SIFa neuron active phases demarcated in light blue shaded region.
**b,c.** Normalized duty cycle, average signal, and active phase duration, and true-valued duty cycle (inset) and active phase duration plotted for the indicated conditions. Shaded areas, s.d. Closed circles mark where the experimental distributions differ from CTRL ($p<0.05$) by two-sample KS test.
**d.** Active phase rise and fall time differences and ratio of active phase durations between SIFa neurons and brainwide PSINA (recorded from SIFa neuron ROI) in CTRL (black) animals or with SIFaR overexpression (red). ~20 data points (range: 8-56) per hourly distribution compiled from each experimental set. Shaded areas, s.d. Closed circles mark where the red distributions differ from black ($p<0.05$) by two-sample KS test.
Both genotypes included the same *GAL80ts* allele (see Supplementary Table S1); rearing and data collection were carried out at 29$^{o}$C to allow for uninhibited expression of the UAS construct.

**Extended Data Figure 7. *NorpA* expression in SIFa neurons rescues major *norpA* mutant phenotype.**
**a.** *Top:* Temperature schedule used. *Bottom:* Representative PSINA traces at 29$^{o}$C for the indicated genotypes.
**b.** Duty cycle, normalized to control, and major period duration for the genotypes and sample sizes in (a). Inset in duty cycle plots true values of metric. Major period corresponds to the dominant frequency in the PSD. Shaded areas, s.d. Closed circles mark where experimental distributions differ from control ($p<0.05$) by two-sample KS test.
**c.** Average PSDs from control and experimental animals; presented as in Fig.1g.

**Extended Data Figure 8. *SIFa*$^{LexA}$ drivers; Related to Fig.5.**

**a.** MIP of 72hAPF brain with *SIFa*$^{LexA.2}$ (i.e., *14F05*$^{LexA}$) driving *myr::SM-V5* (cyan-hot LUT); brain also stained against SIFa (orange-hot LUT) and N-Cadherin (magenta). Yellow square marks location of SIFa soma. Scale bar, 100µm.
**b.** MIP of 72hAPF brain with *SIFa*$^{LexA.3}$ (i.e., *59A06*$^{LexA}$) driving *myr::SM-V5* (stained, cyan-hot LUT); brain also stained against SIFa (orange-hot LUT) and N-Cadherin (magenta). Yellow square marks location of SIFa soma.
**c.** MIP of a 72hAPF brain with *SIFa*$^{LexA.2+.3}$ driving *LexAop-GCaMP6s* (anti-GFP, cyan-hot LUT) and *Trpγ*± driving *UAS-KiR2.1-T2A-tdTOM.HA* (anti-HA, orange-hot LUT). Brain also stained against N-Cadherin (magenta); native fluorescence of *tdTOM* is also visible in this channel. Yellow square marks location of SIFa soma. Yellow asterisks mark trachea, visible due to autofluorescence.

**Supplementary Video 1.** The Widefield Assay: Brainwide PSINA from 68 pupae (8 genotypes, 7-9 replicates each) was recorded for 50 hrs.

**Supplementary Video 2.** 2-channel recording of SIFa neuron activity (blue) and brainwide PSINA (magenta) using the widefield assay; related to Fig.4.

### Extended Data Figure 1

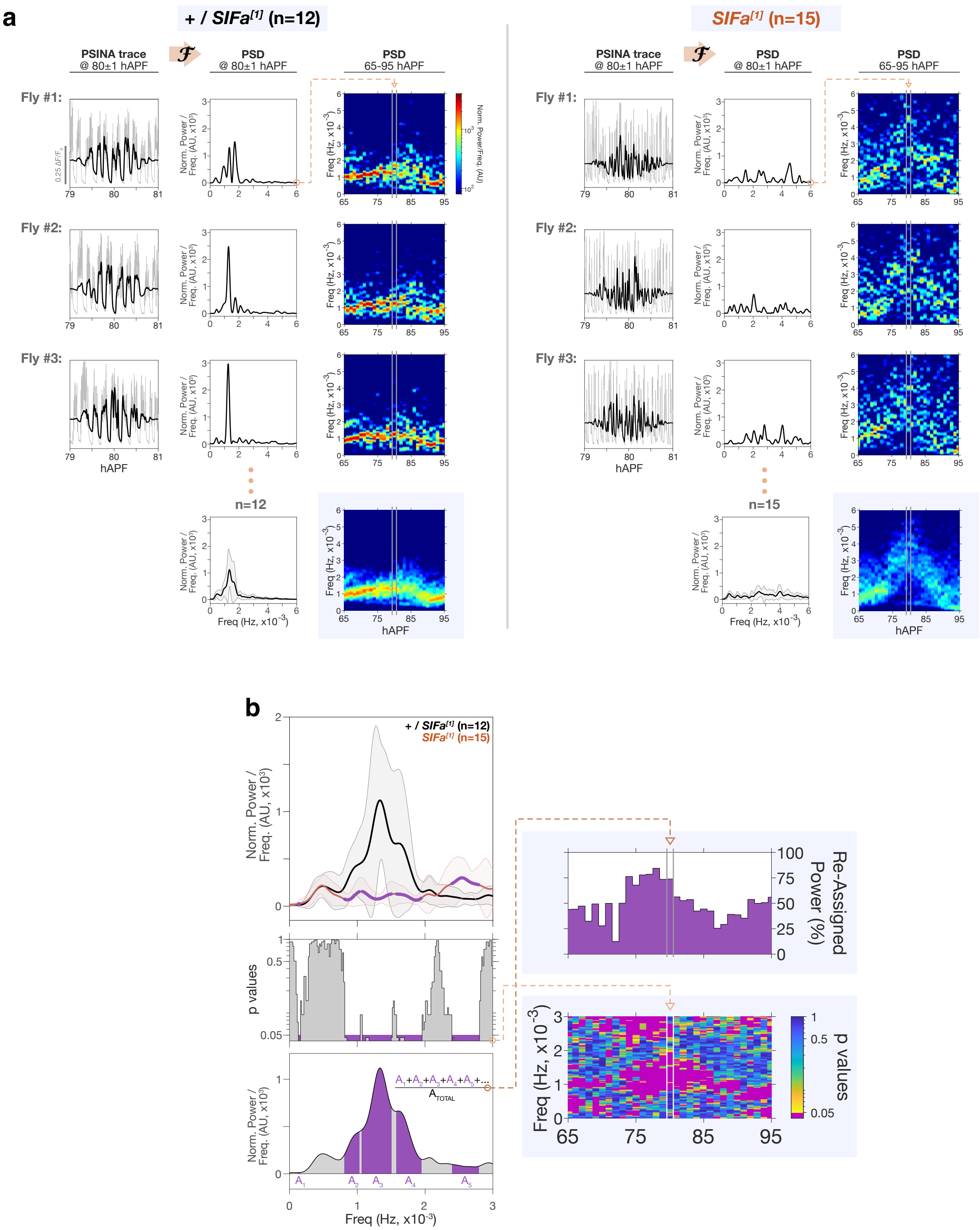

### Extended Data Figure 2

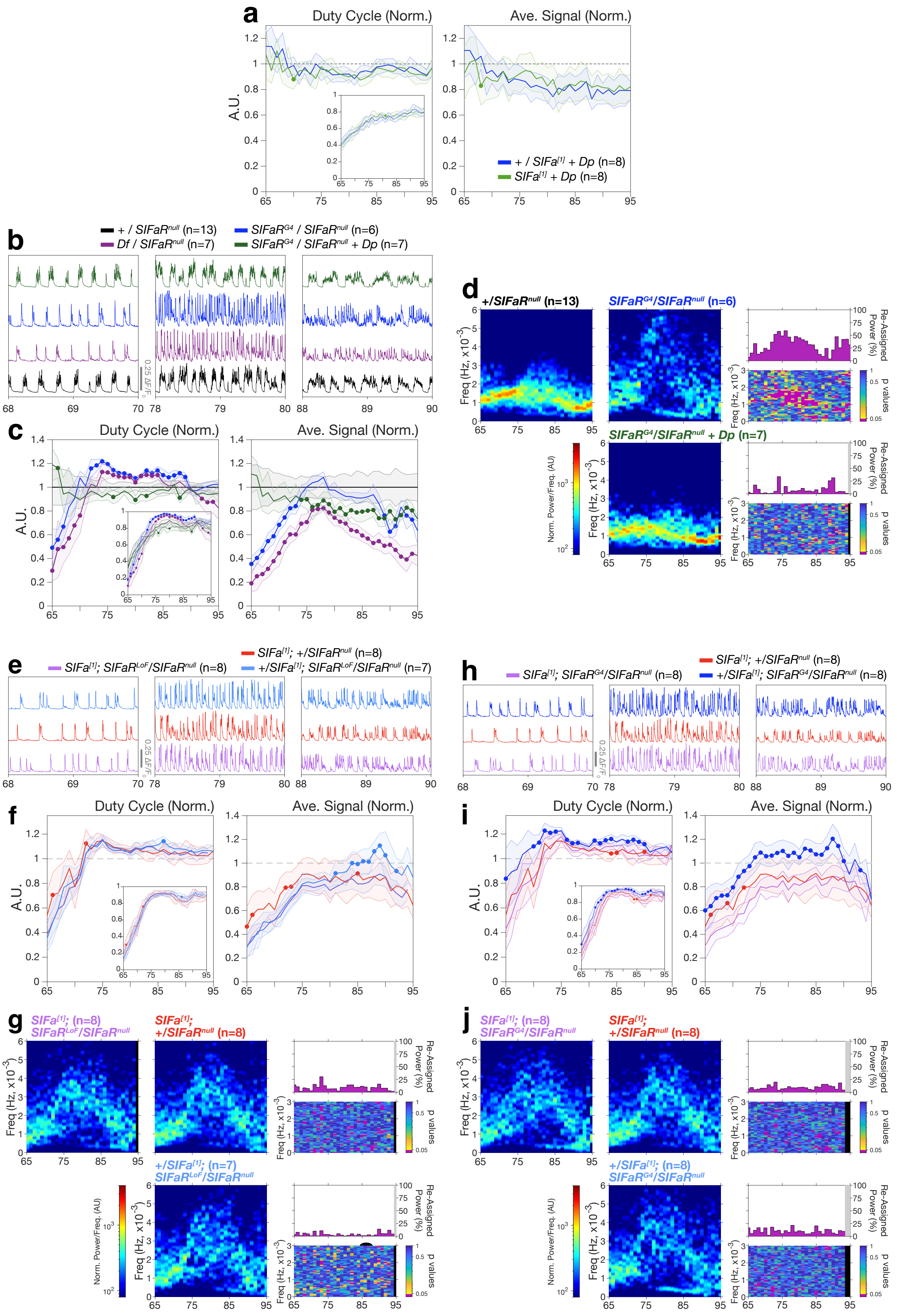

### Extended Data Figure 3

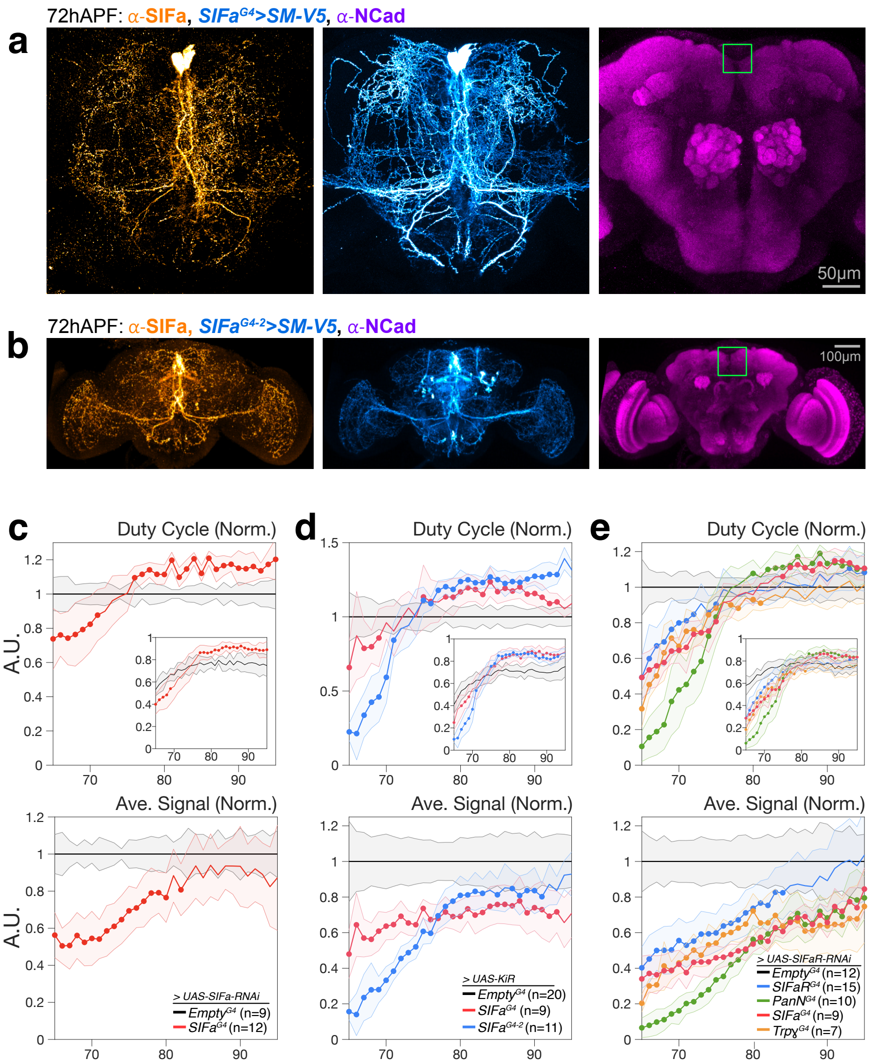

### Extended Data Figure 4

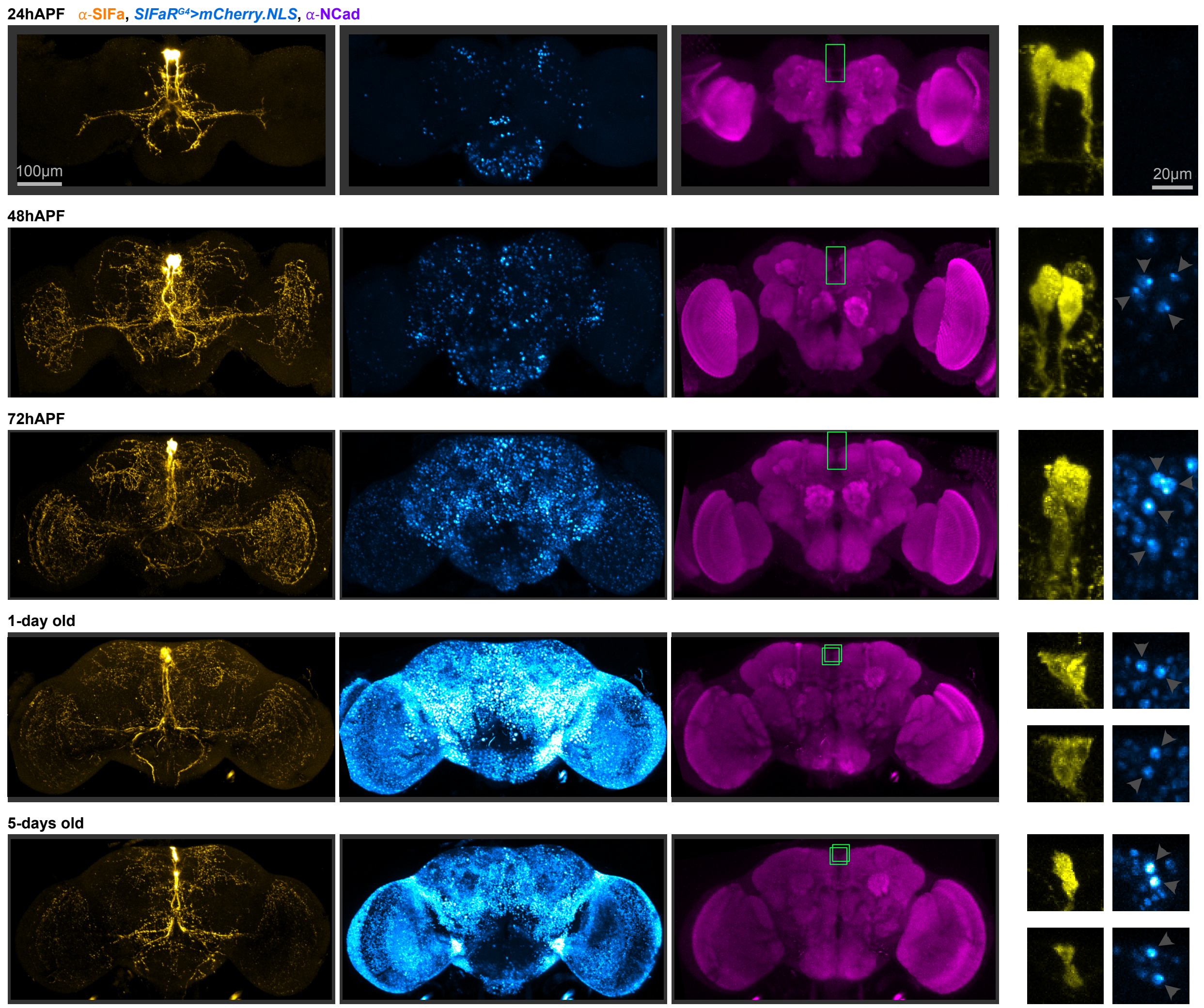

### Extended Data Figure 5

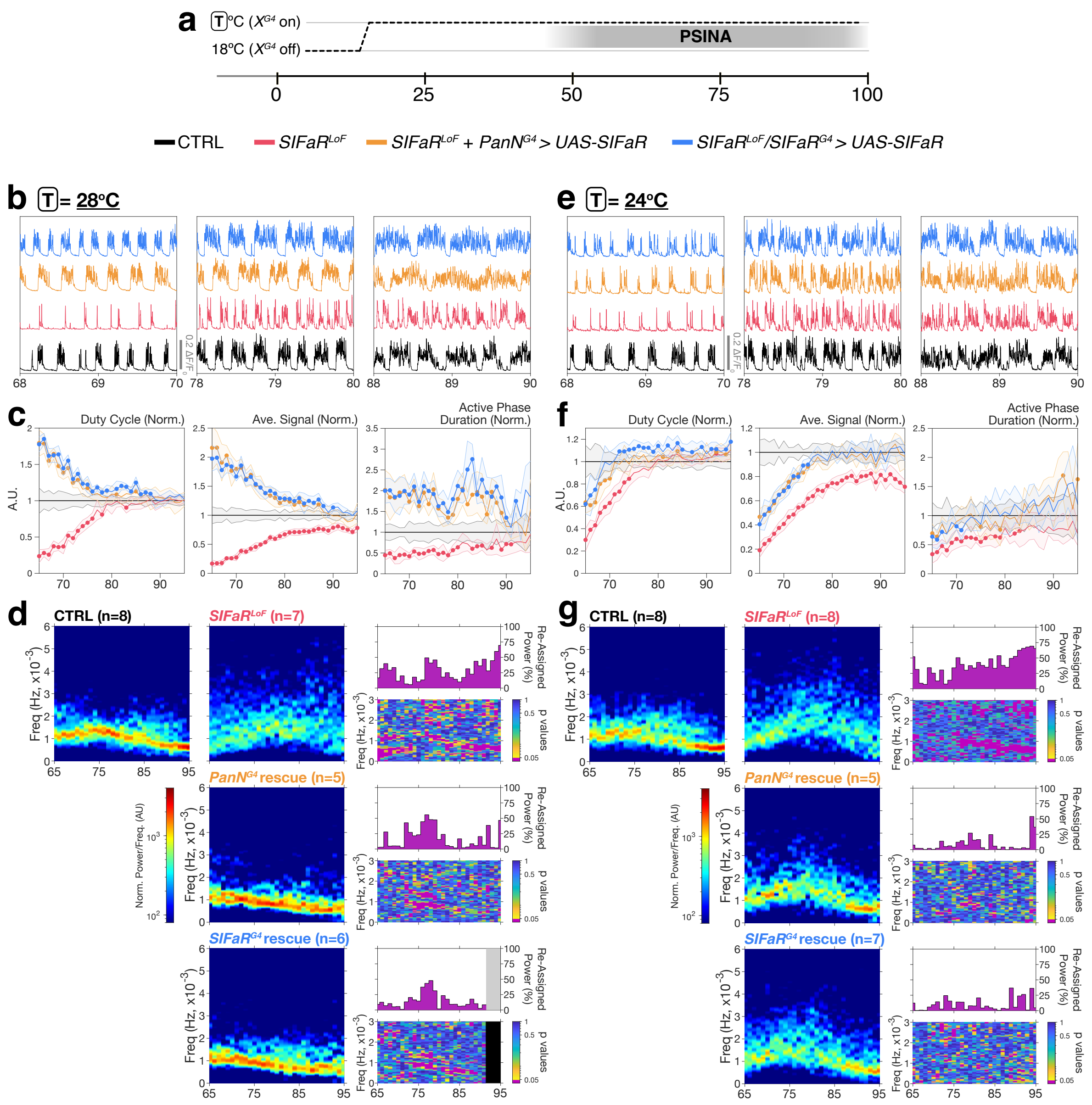

### Extended Data Figure 6

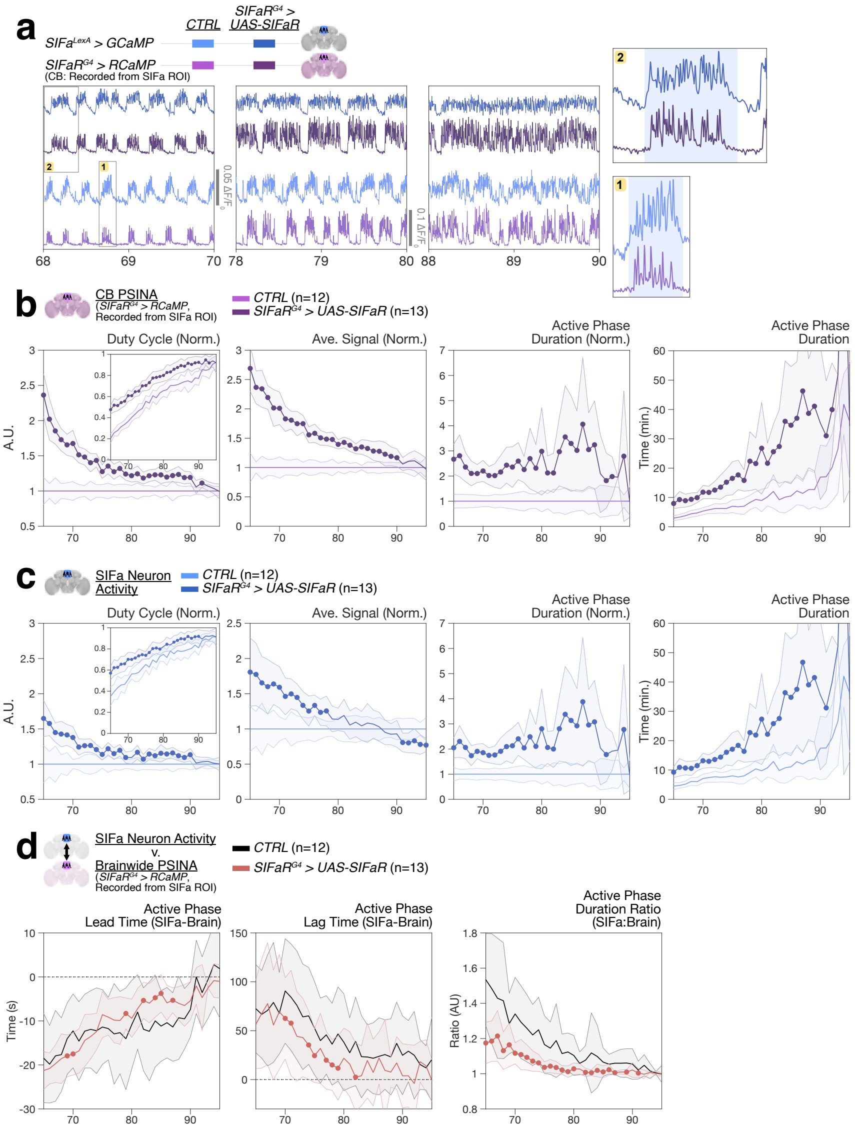

### Extended Data Figure 7

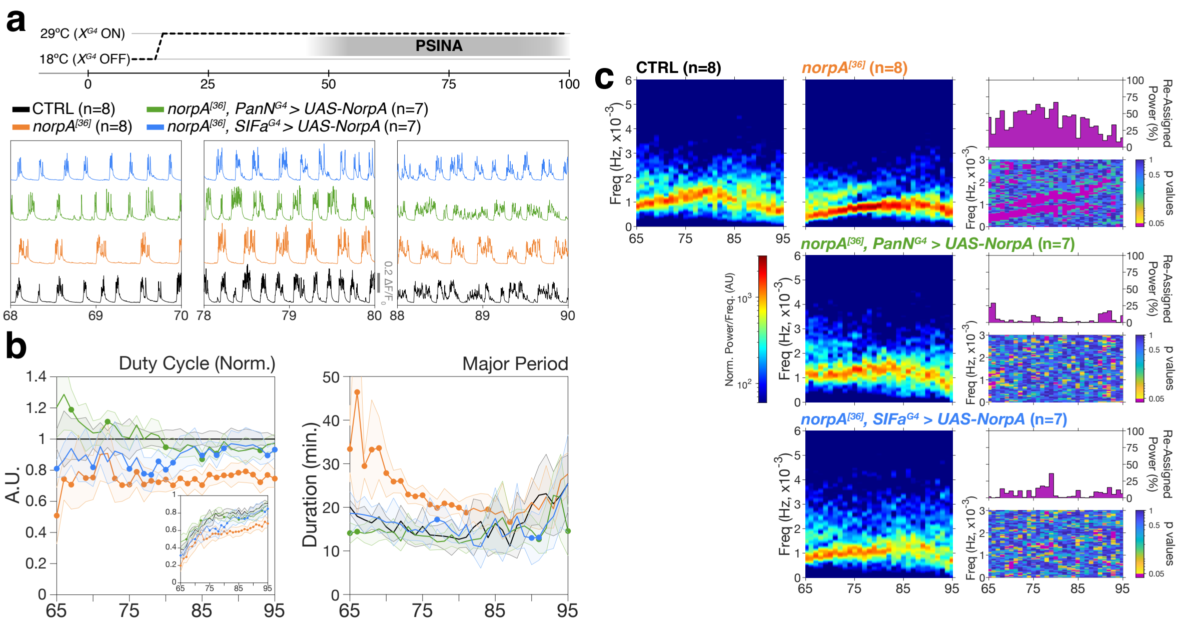

### Extended Data Figure 8

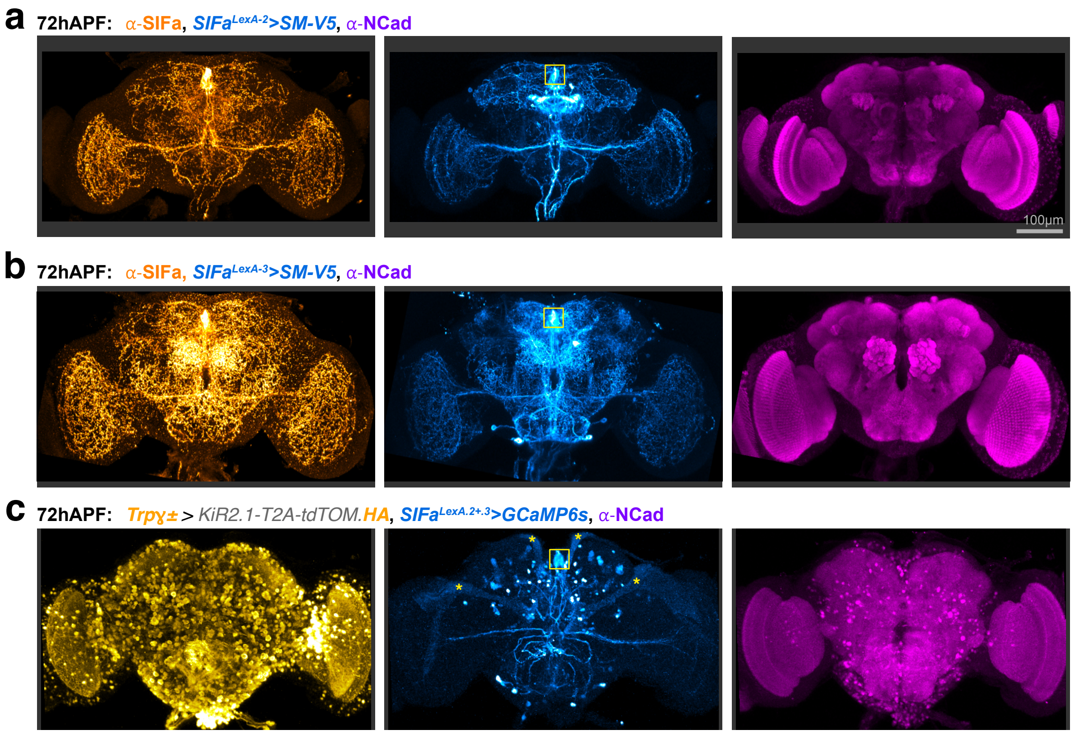

### Supplementary Table 1

| Figure | Label in Figure | Genotype | Notes |
|---|---|---|---|
| **Fig1b-c** | | *w;<br>SIFa[1] / +;<br>13XLexAop2-opGCaMP6s (su(Hw)attP1), R57C10-lexA (VK20)/ +* | |
| **Fig1d** | | *w;<br>Sp-CyO / + ;<br>UAS-mCherry.NLS / SIFaR[MI05376-TG4.1]* | |
| **Fig1e-g** | + / SIFa[1] | *w;<br>SIFa[1] / +;<br>13XLexAop2-opGCaMP6s (su(Hw)attP1), R57C10-lexA (VK20) / +* | |
| | Df / SIFa[1] | *w;<br>SIFa[1] / Df(2R)BSC780;<br>13XLexAop2-opGCaMP6s (su(Hw)attP1), R57C10-lexA (VK20) / +* | |
| | SIFa[1] | *w;<br>SIFa[1];<br>13XLexAop2-opGCaMP6s (su(Hw)attP1), R57C10-lexA (VK20) / +* | |
| | SIFa[2] / SIFa[1] | *w;<br>SIFa[2] / SIFa[1];<br>13XLexAop2-opGCaMP6s (su(Hw)attP1), R57C10-lexA (VK20) / +* | |
| | SIFa[1] + Dp | *w;<br>SIFa[1];<br>13XLexAop2-opGCaMP6s (su(Hw)attP1), R57C10-lexA (VK20) / Dp(2;3)GV-CH321-58A04 (VK31)* | |
| **Fig1h-j** | + / SIFaR null | *w, hs-FLP.D5 (attP3) / w;<br>R57C10-lexA (attP40) , 13XLexAop2-opGCaMP6s (su(Hw)attP5) / +;<br>SIFaR[attP] / +* | FLP element not used in experiments |
| | Df / SIFaR null | *w, hs-FLP.D5 (attP3) / w;<br>R57C10-lexA (attP40) , 13XLexAop2-opGCaMP6s (su(Hw)attP5) / +;<br>SIFaR[attP] / Df(3R)ED10845* | FLP element not used in experiments |
| | SIFaR LoF / SIFaR null | *w, hs-FLP.D5 (attP3) / w;<br>R57C10-lexA (attP40) , 13XLexAop2-opGCaMP6s (su(Hw)attP5) / +;<br>SIFaR[attP] / SIFaR[MI05376-GFSTF.1]* | FLP element not used in experiments |
| | SIFaR LoF / SIFaR null + Dp | *w, hs-FLP.D5 (attP3) / w;<br>R57C10-lexA (attP40) , 13XLexAop2-opGCaMP6s (su(Hw)attP5) / Dp(3;2)GV-CH321-04D21 (VK37);<br>SIFaR[attP] / SIFaR[MI05376-GFSTF.1]* | FLP element not used in experiments |
| **Fig2b,c** | Empty G4 | *UAS-Dcr2, w / w;<br>R57C10-lexA (attP40) , tubP-GAL80[ts](10), 13XLexAop2-opGCaMP6s (su(Hw)attP5) / +;<br>TRiP.JF03364 / empty-GAL4 (attP2)* | |
| | SIFa G4 | *UAS-Dcr2, w / w;<br>R57C10-lexA (attP40) , tubP-GAL80[ts](10), 13XLexAop2-opGCaMP6s (su(Hw)attP5) / SIFa-GAL4;<br>TRiP.JF03364 / +* | |
| **Fig2d,e** | Empty G4 | *yw, UAS-Kir2.1-2A-tdTOM-3xHA (attP3) / w;<br>tubP-GAL80[ts](10) / +;<br>UAS-His-RFP, 13XLexAop2-opGCaMP6s (su(Hw)attP1), R57C10-lexA (VK20) / empty-GAL4 (attP2)* | |
| | SIFa G4 | *yw, UAS-Kir2.1-2A-tdTOM-3xHA (attP3) / w;<br>tubP-GAL80[ts](10) / SIFa-GAL4;<br>UAS-His-RFP, 13XLexAop2-opGCaMP6s (su(Hw)attP1), R57C10-lexA (VK20) / +* | |
| | SIFa G4-2 | *yw, UAS-Kir2.1-2A-tdTOM-3xHA (attP3) / w;<br>tubP-GAL80[ts](10) / +;<br>UAS-His-RFP, 13XLexAop2-opGCaMP6s (su(Hw)attP1), R57C10-lexA (VK20) / R14F05-GAL4 (attP2)* | |
| **Fig2f,g** | Empty G4 | *UAS-Dcr2, w / w;<br>R57C10-lexA (attP40) , tubP-GAL80[ts](10), 13XLexAop2-opGCaMP6s (su(Hw)attP5) / +;<br>TRiP.JF01849 / empty-GAL4 (attP2)* | |
| | SIFaR G4 | *UAS-Dcr2, w / w;<br>R57C10-lexA (attP40) , tubP-GAL80[ts](10), 13XLexAop2-opGCaMP6s (su(Hw)attP5) / Sp-CyO;<br>TRiP.JF01849 / SIFaR[MI05376-TG4.1]* | |
| | PanN G4 | *UAS-Dcr2, w / w;<br>R57C10-lexA (attP40) , tubP-GAL80[ts](10), 13XLexAop2-opGCaMP6s (su(Hw)attP5) / R57C10-GAL4 (JK22C);<br>TRiP.JF01849 / SIFaR[attP]* | |
| | Trpγ G4 | *UAS-Dcr2, w / w;<br>R57C10-lexA (attP40) , tubP-GAL80[ts](10), 13XLexAop2-opGCaMP6s (su(Hw)attP5) / TrpGamma[G4];<br>TRiP.JF01849 / SIFaR[attP]* | |
| | SIFa G4 | *UAS-Dcr2, w / w;<br>R57C10-lexA (attP40) , tubP-GAL80[ts](10), 13XLexAop2-opGCaMP6s (su(Hw)attP5) / SIFa-GAL4;<br>TRiP.JF01849 / SIFaR[attP]* | |
| **Fig2h** | Empty G4 | *yw, UAS-Kir2.1-2A-tdTOM-3xHA (attP3) / w;<br>tubP-GAL80[ts](10) / +;<br>UAS-His-RFP, 13XLexAop2-opGCaMP6s (su(Hw)attP1), R57C10-lexA (VK20) / empty-GAL4 (attP2)* | |
| | SIFaR G4 | *yw, UAS-Kir2.1-2A-tdTOM-3xHA (attP3) / w;<br>tubP-GAL80[ts](10) / Sp-CyO;<br>UAS-His-RFP, 13XLexAop2-opGCaMP6s (su(Hw)attP1), R57C10-lexA (VK20) / SIFaR[MI05376-TG4.1]* | |
| **Fig3b-e** | Empty G4 (CB) | *yw, UAS-Kir2.1-2A-tdTOM-3xHA (attP3) / R29C07-FLP1 (su(Hw)attP8);<br>tubP-GAL80[ts](10), tubP(FRT.stop)GAL80}2 / +;<br>13XLexAop2-opGCaMP6s (su(Hw)attP1), R57C10-lexA (VK20) / empty-GAL4 (attP2)* | |
| | PanN G4 (CB) | *yw, UAS-Kir2.1-2A-tdTOM-3xHA (attP3) / R29C07-FLP1 (su(Hw)attP8);<br>tubP-GAL80[ts](10), tubP(FRT.stop)GAL80}2 / +;<br>13XLexAop2-opGCaMP6s (su(Hw)attP1), R57C10-lexA (VK20) / R57C10-GAL4 (attP2)* | |
| | Trpγ G4 (CB) | *yw, UAS-Kir2.1-2A-tdTOM-3xHA (attP3) / R29C07-FLP1 (su(Hw)attP8);<br>tubP-GAL80[ts](10), tubP(FRT.stop)GAL80}2 / TrpGamma[G4];<br>13XLexAop2-opGCaMP6s (su(Hw)attP1), R57C10-lexA (VK20) / +* | |
| | SIFaR G4 (CB) | *yw, UAS-Kir2.1-2A-tdTOM-3xHA (attP3) / R29C07-FLP1 (su(Hw)attP8);<br>tubP-GAL80[ts](10), tubP(FRT.stop)GAL80}2 / +;<br>13XLexAop2-opGCaMP6s (su(Hw)attP1), R57C10-lexA (VK20) / SIFaR[MI05376-TG4.1]* | |
| | Empty G4 (OL) | *yw, UAS-Kir2.1-2A-tdTOM-3xHA (attP3) / R29C07-FLP1 (su(Hw)attP8);<br>tubP-GAL80[ts](10), alphaTub84B(FRT.GAL80)}2 / +;<br>13XLexAop2-opGCaMP6s (su(Hw)attP1), R57C10-lexA (VK20) / empty-GAL4 (attP2)* | |
| | PanN G4 (OL) | *yw, UAS-Kir2.1-2A-tdTOM-3xHA (attP3) / R29C07-FLP1 (su(Hw)attP8);<br>tubP-GAL80[ts](10), alphaTub84B(FRT.GAL80)}2 / +;<br>13XLexAop2-opGCaMP6s (su(Hw)attP1), R57C10-lexA (VK20) / R57C10-GAL4 (attP2)* | |
| | Trpγ G4 (OL) | *yw, UAS-Kir2.1-2A-tdTOM-3xHA (attP3) / R29C07-FLP1 (su(Hw)attP8);<br>tubP-GAL80[ts](10), alphaTub84B(FRT.GAL80)}2 / TrpGamma[G4];<br>13XLexAop2-opGCaMP6s (su(Hw)attP1), R57C10-lexA (VK20) / +* | |
| | SIFaR G4 (OL) | *yw, UAS-Kir2.1-2A-tdTOM-3xHA (attP3) / R29C07-FLP1 (su(Hw)attP8);<br>tubP-GAL80[ts](10), alphaTub84B(FRT.GAL80)}2 / +;<br>13XLexAop2-opGCaMP6s (su(Hw)attP1), R57C10-lexA (VK20) / SIFaR[MI05376-TG4.1]* | |
| **Fig3f-h** | CTRL (CB) | *UAS-Dcr2, w / R29C07-FLP1 (su(Hw)attP8);<br>tubP-GAL80[ts](10), tubP(FRT.stop)GAL80}2 / +;<br>13XLexAop2-opGCaMP6s (su(Hw)attP1), SIFaR[attP], R57C10-lexA (VK20) / TRiP.JF01849* | |
| | PanN G4 (CB) | *UAS-Dcr2, w / R29C07-FLP1 (su(Hw)attP8);<br>tubP-GAL80[ts](10), tubP(FRT.stop)GAL80}2 / R57C10-GAL4 (JK22C);<br>13XLexAop2-opGCaMP6s (su(Hw)attP1), SIFaR[attP], R57C10-lexA (VK20) / TRiP.JF01849* | |
| | CTRL (OL) | *UAS-Dcr2, w / R29C07-FLP1 (su(Hw)attP8);<br>tubP-GAL80[ts](10), alphaTub84B(FRT.GAL80)}2 / +;<br>13XLexAop2-opGCaMP6s (su(Hw)attP1), SIFaR[attP], R57C10-lexA (VK20) / TRiP.JF01849* | |
| | PanN G4 (OL) | *UAS-Dcr2, w / R29C07-FLP1 (su(Hw)attP8);<br>tubP-GAL80[ts](10), alphaTub84B(FRT.GAL80)}2 / R57C10-GAL4 (JK22C);<br>13XLexAop2-opGCaMP6s (su(Hw)attP1), SIFaR[attP], R57C10-lexA (VK20) / TRiP.JF01849* | |
| **Fig4a-g** | CTRL | *13XLexAop2-opGCaMP6s (?, su(Hw)attp8) / w;<br>SIFa[1] / +;<br>R57C10-GAL4(attP2), 20XUAS-IVS-NES-jRCaMP1b-p10 (VK5) / SIFa-LexA.GAD* | Recombinant of two independent 13XLexAop2-opGCaMP6s insertions |
| | SIFa null | *13XLexAop2-opGCaMP6s (?, su(Hw)attp8) / w;<br>SIFa[1];<br>R57C10-GAL4(attP2), 20XUAS-IVS-NES-jRCaMP1b-p10 (VK5) / SIFa-LexA.GAD* | Recombinant of two independent 13XLexAop2-opGCaMP6s insertions |
| **Fig4h,i** | CTRL | *13XLexAop2-opGCaMP6s (?, su(Hw)attp8) /w;<br>SIFa[1] / +;<br>R57C10-GAL4(attP2), 20XUAS-IVS-NES-jRCaMP1b-p10 (VK5) / SIFa-LexA.GAD* | Recombinant of two independent 13XLexAop2-opGCaMP6s insertions |
| | TrpA1 | *13XLexAop2-opGCaMP6s (?, su(Hw)attp8) /w;<br>SIFa[1] / 13XLexAop2-IVS-dTrpA1-WPRE (su(Hw)attP5);<br>R57C10-GAL4(attP2), 20XUAS-IVS-NES-jRCaMP1b-p10 (VK5) / SIFa-LexA.GAD* | Recombinant of two independent 13XLexAop2-opGCaMP6s insertions |

| Figure | Label in Figure | Genotype | Notes |
|---|---|---|---|
| Fig4j-l | Het, CTRL | *13XLexAop2-opGCaMP6s (?, su(Hw)attp8) / w;*<br>*SIFa[1] / +;*<br>*R57C10-GAL4(attP2), 20XUAS-IVS-NES-jRCaMP1b-p10 (VK5) / SIFa-LexA.GAD* | Recombinant of two independent 13XLexAop2-opGCaMP6s insertions |
|  | Het, TrpA1 | *13XLexAop2-opGCaMP6s (?, su(Hw)attp8) / w;*<br>*SIFa[1] / 13XLexAop2-IVS-dTrpA1-WPRE (su(Hw)attP5);*<br>*R57C10-GAL4(attP2), 20XUAS-IVS-NES-jRCaMP1b-p10 (VK5) / SIFa-LexA.GAD* | Recombinant of two independent 13XLexAop2-opGCaMP6s insertions |
|  | SIFa null, CTRL | *13XLexAop2-opGCaMP6s (?, su(Hw)attp8) / w;*<br>*SIFa[1];*<br>*R57C10-GAL4(attP2), 20XUAS-IVS-NES-jRCaMP1b-p10 (VK5) / SIFa-LexA.GAD* | Recombinant of two independent 13XLexAop2-opGCaMP6s insertions |
|  | SIFa null, TrpA1 | *13XLexAop2-opGCaMP6s (?, su(Hw)attp8) / w;*<br>*SIFa[1] / 13XLexAop2-IVS-dTrpA1-WPRE (su(Hw)attP5), SIFa[1];*<br>*R57C10-GAL4(attP2), 20XUAS-IVS-NES-jRCaMP1b-p10 (VK5) / SIFa-LexA.GAD* | Recombinant of two independent 13XLexAop2-opGCaMP6s insertions |
| Fig5b-e | CTRL | *13XLexAop2-opGCaMP6s (?, su(Hw)attp8) / w;*<br>*R14F05-lexA (attP40) , Trpɣ[CR70048-TG4.2] / R59A06-LexA (attP40);*<br>*8XLexAop2-IVS-GAL80-WPRE (attP2) / +* | Recombinant of two independent 13XLexAop2-opGCaMP6s insertions |
| | Trpɣ± > KiR2.1 | *13XLexAop2-opGCaMP6s (?, su(Hw)attp8) / yw, UAS-Kir2.1-2A-tdTOM-3xHA (attP3) ;*<br>*R14F05-lexA (attP40) , Trpɣ[CR70048-TG4.2] / R59A06-LexA (attP40);*<br>*8XLexAop2-IVS-GAL80-WPRE (attP2) / +* | |
| Fig6a-c | CTRL | *w;*<br>*UAS-SIFaR / +;*<br>*13XLexAop2-opGCaMP6s (su(Hw)attP1), SIFaR[attP], R57C10-lexA (VK20) / empty-GAL4 (attP2)* |  |
|  | OE: SIFa G4>UAS-SIFaR | *w;*<br>*UAS-SIFaR / SIFa-GAL4;*<br>*13XLexAop2-opGCaMP6s (su(Hw)attP1), SIFaR[attP], R57C10-lexA (VK20) / +* |  |
| | SIFaR LoF/ SIFaR null | *w;*<br>*UAS-SIFaR / +;*<br>*13XLexAop2-opGCaMP6s (su(Hw)attP1), SIFaR[attP], R57C10-lexA (VK20) / SIFaR[MI05376-GFSTF.1]* | |
| | + SIFa G4>UAS-SIFaR | *w;*<br>*UAS-SIFaR / SIFa-GAL4;*<br>*13XLexAop2-opGCaMP6s (su(Hw)attP1), SIFaR[attP], R57C10-lexA (VK20) / SIFaR[MI05376-GFSTF.1]* | |
| | +Trpɣ G4 >UAS-SIFaR | *w;*<br>*UAS-SIFaR / TrpGamma[G4];*<br>*13XLexAop2-opGCaMP6s (su(Hw)attP1), SIFaR[attP], R57C10-lexA (VK20) / SIFaR[MI05376-GFSTF.1]* | |
| | +Trpɣ G4.2 >2x UAS-SIFaR | *w;*<br>*UAS-SIFaR / UAS-SIFaR, Trpɣ[CR70048-TG4.2];*<br>*13XLexAop2-opGCaMP6s (su(Hw)attP1), SIFaR[attP], R57C10-lexA (VK20) / SIFaR[MI05376-GFSTF.1]* | |
| Fig7a-c | CTRL | *w, 20XUAS-IVS-opGCaMP6s (su(Hw)attP8) / w;*<br>*R57C10-lexA (attP40) , 13XLexAop2-jRCaMP1b (su(Hw)attP5), SIFa[1] / Trpɣ[CR70048-TG4.2];*<br>*20XUAS-IVS-opGCaMP6s (su(Hw)attP1) / +* | Fluorescent eye marker was removed from Trpɣ[CR70048-TG4.2] using Cre |
| | SIFa null | *w, 20XUAS-IVS-opGCaMP6s (su(Hw)attP8) / w;*<br>*R57C10-lexA (attP40) , 13XLexAop2-jRCaMP1b (su(Hw)attP5), SIFa[1] / Trpɣ[CR70048-TG4.2], SIFa[2];*<br>*20XUAS-IVS-opGCaMP6s (su(Hw)attP1) / +* | Fluorescent eye marker was removed from Trpɣ[CR70048-TG4.2] using Cre |
| FigED1a,b | + / SIFa[1] | *w;*<br>*SIFa[1] / +;*<br>*13XLexAop2-opGCaMP6s (su(Hw)attP1), R57C10-lexA (VK20) / +* | |
| | SIFa[1] | *w;*<br>*SIFa[1];*<br>*13XLexAop2-opGCaMP6s (su(Hw)attP1), R57C10-lexA (VK20) / +* | |
| FigED2a | + / SIFa[1] + Dp | *w;*<br>*SIFa[1] / Sp - CyO - +;*<br>*13XLexAop2-opGCaMP6s (su(Hw)attP1), R57C10-lexA (VK20) / Dp(2;3)GV-CH321-58A04 (VK31)* | |
| | SIFa[1] + Dp | *w;*<br>*SIFa[1];*<br>*13XLexAop2-opGCaMP6s (su(Hw)attP1), R57C10-lexA (VK20) / Dp(2;3)GV-CH321-58A04 (VK31)* | Same as in Fig1e-g |
| FigED2b-d | + / SIFaR null | *w, hs-FLP.D5 (attP3) / w;*<br>*R57C10-lexA (attP40) , 13XLexAop2-opGCaMP6s (su(Hw)attP5) / +;*<br>*SIFaR[attP] / +* | FLP element not used in experiments |
| | Df / SIFaR null | *w, hs-FLP.D5 (attP3) / w;*<br>*R57C10-lexA (attP40) , 13XLexAop2-opGCaMP6s (su(Hw)attP5) / +;*<br>*SIFaR[attP] / Df(3R)ED10845* | FLP element not used in experiments |
| | SIFaR G4 / SIFaR null | *w, hs-FLP.D5 (attP3) / w;*<br>*R57C10-lexA (attP40) , 13XLexAop2-opGCaMP6s (su(Hw)attP5) / +;*<br>*SIFaR[attP] / SIFaR[MI05376-TG4.1]* | FLP element not used in experiments |
| | SIFaR G4 / SIFaR null + Dp | *w, hs-FLP.D5 (attP3) / w;*<br>*R57C10-lexA (attP40) , 13XLexAop2-opGCaMP6s (su(Hw)attP5) / Dp(3;2)GV-CH321-04D21 (VK37);*<br>*SIFaR[attP] / SIFaR[MI05376-TG4.1]* | FLP element not used in experiments |
| FigED2e-g | SIFa[1]; SIFaR LoF / SIFaR null | *w, hs-FLP.D5 (attP3) / w;*<br>*SIFa[1];*<br>*13XLexAop2-opGCaMP6s (su(Hw)attP1), SIFaR[attP], R57C10-lexA (VK20) / SIFaR[MI05376-GFSTF.1]* | FLP element not used in experiments |
| | SIFa[1]; + / SIFaR null | *w, hs-FLP.D5 (attP3) / w;*<br>*SIFa[1];*<br>*13XLexAop2-opGCaMP6s (su(Hw)attP1), SIFaR[attP], R57C10-lexA (VK20) / +* | FLP element not used in experiments |
| | + / SIFa[1]; SIFaR LoF / SIFaR nul | *w, hs-FLP.D5 (attP3) / w;*<br>*SIFa[1] / +;*<br>*13XLexAop2-opGCaMP6s (su(Hw)attP1), SIFaR[attP], R57C10-lexA (VK20) / SIFaR[MI05376-GFSTF.1]* | FLP element not used in experiments |
| FigED2h-j | SIFa[1]; SIFaR G4 / SIFaR null | *w, hs-FLP.D5 (attP3) / w;*<br>*SIFa[1];*<br>*13XLexAop2-opGCaMP6s (su(Hw)attP1), SIFaR[attP], R57C10-lexA (VK20) / SIFaR[MI05376-TG4.1]* | FLP element not used in experiments |
| | SIFa[1]; + / SIFaR null | *w, hs-FLP.D5 (attP3) / w;*<br>*SIFa[1];*<br>*13XLexAop2-opGCaMP6s (su(Hw)attP1), SIFaR[attP], R57C10-lexA (VK20) / +* | FLP element not used in experiments |
| | + / SIFa[1]; SIFaR G4 / SIFaR null | *w, hs-FLP.D5 (attP3) / w;*<br>*SIFa[1] / +;*<br>*13XLexAop2-opGCaMP6s (su(Hw)attP1), SIFaR[attP], R57C10-lexA (VK20) / SIFaR[MI05376-TG4.1]* | FLP element not used in experiments |
| FigED3a | | *w, 10XUAS-IVS-myr::smGdP-V5 (attP18) / w;*<br>*SIFa-GAL4 / ap[Xa];*<br>*Sb / +* | |
| FigED3b | | *w, 10XUAS-IVS-myr::smGdP-V5 (attP18) / w;*<br>*+/ Sp;*<br>*R14F05-GAL4 (attP2) / +* | |
| FigED3c | Empty G4 | *UAS-Dcr2, w / w;*<br>*R57C10-lexA (attP40) , tubP-GAL80[ts](10), 13XLexAop2-opGCaMP6s (su(Hw)attP5) / +;*<br>*TRiP.JF03364 / empty-GAL4 (attP2)* | Same as in Fig2b,c |
| | SIFa G4 | *UAS-Dcr2, w / w;*<br>*R57C10-lexA (attP40) , tubP-GAL80[ts](10), 13XLexAop2-opGCaMP6s (su(Hw)attP5) / SIFa-GAL4;*<br>*TRiP.JF03364 / +* | Same as in Fig2b,c |
| FigED3d | Empty G4 | *yw, UAS-Kir2.1-2A-tdTOM-3xHA (attP3) / w;*<br>*tubP-GAL80[ts](10) / +;*<br>*UAS-His-RFP, 13XLexAop2-opGCaMP6s (su(Hw)attP1), R57C10-lexA (VK20) / empty-GAL4 (attP2)* | Same as in Fig2d,e |
|  | SIFa G4 | *yw, UAS-Kir2.1-2A-tdTOM-3xHA (attP3) / w;*<br>*tubP-GAL80[ts](10) / SIFa-GAL4;*<br>*UAS-His-RFP, 13XLexAop2-opGCaMP6s (su(Hw)attP1), R57C10-lexA (VK20) / +* | Same as in Fig2d,e |
|  | SIFa G4-2 | *yw, UAS-Kir2.1-2A-tdTOM-3xHA (attP3) / w;*<br>*tubP-GAL80[ts](10) / +;*<br>*UAS-His-RFP, 13XLexAop2-opGCaMP6s (su(Hw)attP1), R57C10-lexA (VK20) / R14F05-GAL4 (attP2)* | Same as in Fig2d,e |
| FigED3e | Empty G4 | *UAS-Dcr2, w / w;*<br>*R57C10-lexA (attP40) , tubP-GAL80[ts](10), 13XLexAop2-opGCaMP6s (su(Hw)attP5) / +;*<br>*TRiP.JF01849 / empty-GAL4 (attP2)* | Same as in Fig2f,g |
|  | SIFaR G4 | *UAS-Dcr2, w / w;*<br>*R57C10-lexA (attP40) , tubP-GAL80[ts](10), 13XLexAop2-opGCaMP6s (su(Hw)attP5) / Sp-CyO;*<br>*TRiP.JF01849 / SIFaR[MI05376-TG4.1]* | Same as in Fig2f,g |
|  | PanN G4 | *UAS-Dcr2, w / w;*<br>*R57C10-lexA (attP40) , tubP-GAL80[ts](10), 13XLexAop2-opGCaMP6s (su(Hw)attP5) / R57C10-GAL4 (JK22C);*<br>*TRiP.JF01849 / SIFaR[attP]* | Same as in Fig2f,g |
|  | Trpɣ G4 | *UAS-Dcr2, w / w;*<br>*R57C10-lexA (attP40) , tubP-GAL80[ts](10), 13XLexAop2-opGCaMP6s (su(Hw)attP5) / TrpGamma[G4];*<br>*TRiP.JF01849 / SIFaR[attP]* | Same as in Fig2f,g |
|  | SIFa G4 | *UAS-Dcr2, w / w;*<br>*R57C10-lexA (attP40) , tubP-GAL80[ts](10), 13XLexAop2-opGCaMP6s (su(Hw)attP5) / SIFa-GAL4;*<br>*TRiP.JF01849 / SIFaR[attP]* | Same as in Fig2f,g |

| Figure | Label in Figure | Genotype | Notes |
|---|---|---|---|
| **FigED4** | | *w;*<br>*Sp-CyO / + ;*<br>*UAS-mCherry.NLS / SIFaR[MI05376-TG4.1]* | Same as in Fig1d |
| **FigED5** | CTRL | *w, 20XUAS-IVS-opGCaMP6s (su(Hw)attP8) / +;*<br>*R57C10-lexA (attP40) , 13XLexAop2-jRCaMP1b (su(Hw)attP5), UAS-SIFaR / tubP-GAL80[ts](10);*<br>*SIFaR[MI05376-GFSTF.1] / +* | Only RCaMP channel recorded |
| | SIFaR LoF | *w, 20XUAS-IVS-opGCaMP6s (su(Hw)attP8) / +;*<br>*R57C10-lexA (attP40) , 13XLexAop2-jRCaMP1b (su(Hw)attP5), UAS-SIFaR / +;*<br>*SIFaR[MI05376-GFSTF.1] / +* | Only RCaMP channel recorded |
| | PanN G4 rescue | *w, 20XUAS-IVS-opGCaMP6s (su(Hw)attP8) / +;*<br>*R57C10-lexA (attP40) , 13XLexAop2-jRCaMP1b (su(Hw)attP5), UAS-SIFaR / tubP-GAL80[ts](10);*<br>*SIFaR[MI05376-GFSTF.1] / R57C10-GAL4 (attP2), SIFaR[MI05376-GFSTF.1]* | Only RCaMP channel recorded |
| | SIFaR G4 rescue | *w, 20XUAS-IVS-opGCaMP6s (su(Hw)attP8) / +;*<br>*R57C10-lexA (attP40) , 13XLexAop2-jRCaMP1b (su(Hw)attP5), UAS-SIFaR / tubP-GAL80[ts](10);*<br>*SIFaR[MI05376-GFSTF.1] / SIFaR[MI05376-TG4.1]* | Only RCaMP channel recorded |
| **FigED6** | CTRL | *13XLexAop2-opGCaMP6s (?, su(Hw)attp8) / w;*<br>*tubP-GAL80[ts](10) / +;*<br>*20XUAS-IVS-NES-jRCaMP1b-p10 (VK5), SIFa-LexA.GAD / SIFaR[MI05376-TG4.1]* | Recombinant of two independent 13XLexAop2-opGCaMP6s insertions<br>Crosses and progeny reared at 29oC throughout development |
|  | SIFaR G4 > UAS-SIFaR | *13XLexAop2-opGCaMP6s (?, su(Hw)attp8) / w;*<br>*tubP-GAL80[ts](10) / UAS-SIFaR;*<br>*20XUAS-IVS-NES-jRCaMP1b-p10 (VK5), SIFa-LexA.GAD / SIFaR[MI05376-TG4.1]* | Recombinant of two independent 13XLexAop2-opGCaMP6s insertions<br>Crosses and progeny reared at 29oC throughout development |
| **FigED7** | CTRL | *w, norpA[36] / w;*<br>*SIFa[1] / +;*<br>*13XLexAop2-opGCaMP6s (su(Hw)attP1), R57C10-lexA (VK20), tubP-GAL80[ts](7)/ +* | |
| | norpA[36] | *w, norpA[36];*<br>*SIFa[1] / +;*<br>*13XLexAop2-opGCaMP6s (su(Hw)attP1), R57C10-lexA (VK20), tubP-GAL80[ts](7)/ +* | |
| | norpA[36], PanN G4 | *w, norpA[36];*<br>*SIFa[1] / R57C10-GAL4 (JK22C);*<br>*13XLexAop2-opGCaMP6s (su(Hw)attP1), R57C10-lexA (VK20), tubP-GAL80[ts](7)/ UAS-norpA.K* | |
| | norpA[36], SIFa G4 | *w, norpA[36];*<br>*SIFa[1] / SIFa-GAL4;*<br>*13XLexAop2-opGCaMP6s (su(Hw)attP1), R57C10-lexA (VK20), tubP-GAL80[ts](7)/ UAS-norpA.K* | |
| **FigED8a** | SIFa LexA.2 | *w, 10XUAS-IVS-myr::smGdP-HA (attP18), 13XLexAop2-IVS-myr::smGdP-V5 (su(Hw)attP8) / w;*<br>*R14F05-LexA (attP40) / Sp ;*<br>*+* | UAS element not used in experiment |
| **FigED8b** | SIFa LexA.3 | *w, 10XUAS-IVS-myr::smGdP-HA (attP18), 13XLexAop2-IVS-myr::smGdP-V5 (su(Hw)attP8) / w;*<br>*R59A06-LexA (attP40) / Sp ;*<br>*+* | UAS element not used in experiment |
| **FigED8c** | SIFa LexA.2+3 | *13XLexAop2-opGCaMP6s (?, su(Hw)attp8) / yw, UAS-Kir2.1-2A-tdTOM-3xHA (attP3) ;*<br>*R14F05-lexA (attP40) , Trpγ[CR70048-TG4.2] / R59A06-LexA (attP40);*<br>*8XLexAop2-IVS-GAL80-WPRE (attP2) / +* | Recombinant of two independent 13XLexAop2-opGCaMP6s insertions<br>Same as in Fig5b-e |
